# Soft selection affects introgression dynamics and the viability of populations experiencing intrusion of maladapted individuals

**DOI:** 10.1101/2023.09.25.559312

**Authors:** Thomas Eric Reed, Adam Kane, Philip McGinnity, Ronan James O’Sullivan

**Author notes:** These authors contributed equally (Thomas Eric Reed, Ronan James O’Sullivan).

## Abstract

The deliberate release of captive-bred individuals, the accidental escape of domesticated strains, or the invasion of closely related conspecifics into wild populations can all lead to introgressive hybridisation, which poses a challenge for conservation and wildlife management. Rates of introgression and the magnitude of associated demographic impacts vary widely across ecological contexts. However, the reasons for this variation remain poorly understood. One rarely considered phenomenon in this context is soft selection, wherein relative trait values determine success in intraspecific competition for a limiting resource. Here we develop an eco-genetic model explicitly focussed on understanding the influence soft selection has on the eco-evolutionary dynamics of wild populations experiencing intrusion from foreign/domesticated individuals. While based on a generalised salmonine lifecycle, the model is applicable to any taxon that experiences incursion from locally maladapted genotypes, in addition to phenotype-dependent competition for a limiting resource (e.g., breeding sites, feeding territories). The effects of both acute and chronic intrusion depended strongly on the relative competitiveness of intruders versus locals. When intruders were competitively inferior, soft selection limited their reproductive success (ability to compete for limited spawning sites), which prevented strong introgression or population declines from occurring. In contrast, when intruders were competitively superior, this amplified introgression and led to increased maladaptation of the admixed population. This had negative consequences for population size and population viability. The results were sensitive to the intrusion level, the magnitude of reproductive excess, trait heritability, and the extent to which intruders were maladapted relative to locals. Our findings draw attention to under- appreciated interactions between soft selection and maladaptive hybridisation, which may be critical to determining the impact captive breeding programmes and domesticated escapees can have on otherwise self-sustaining wild populations.

## Introduction

Free-living populations of animals and plants are typically, though not always, reasonably well-adapted to their local environments (Hendry and Gonzalez 2008). Local adaptation can be disrupted, however, by various anthropogenic stressors, which induce maladaptation by affecting the abiotic/biotic selective landscapes (e.g., climate change, species introductions), and/or by shifting trait distributions relative to trait optima (Chevin et al. 2010). Human-mediated hybridisation, for example, can pull populations away from local adaptive peaks (extrinsic outbreeding depression) or lead to a breakdown of adaptive linkage disequilibrium (intrinsic outbreeding depression - (Grabenstein and Taylor 2018)). The scope for introgressive hybridisation has increased in the Anthropocene, as taxa shift their distributions in response to climate change, intentional introductions/translocations occur, and domesticated individuals escape into wild populations (Brennan et al. 2015; Wayne and Shaffer 2016). A major challenge for conservation biology is, thus, to anticipate and respond appropriately to such changes (Kinnison and Hairston 2007).

The release of captive-reared individuals has long been used as a conservation strategy to replenish beleaguered populations (Seddon et al. 2007; Fraser 2008), as well as a wildlife management tool to increase the number of individuals available for harvest (Barbanera et al. 2010; Claussen and Philipp 2022). However, supplemental stocking often fails to provide the desired “demographic boost” to populations that are already naturally self-sustaining and, in some scenarios, can lead to genetic homogenisation (Skaala et al. 2016; Karlsson et al. 2016) and reduced fitness of hatchery individuals and their hybrids in wild environments (Araki et al. 2008; O’Sullivan et al. 2020). For example, stocking of British rivers with hatchery-produced Atlantic salmon (*Salmo salar*) did not on the average improve, and in some cases apparently negatively affected, rod catches (Young 2013). Nevertheless, the practice remains widespread among salmonines (salmon, trout, charr), in particular Pacific salmonids (*Oncorhynchus* spp.), where industrial-scale hatchery programmes exist for the purposes of enhancing fisheries or augmenting endangered populations (Naish et al. 2007). The reduced fitness of captive-reared individuals in the wild is likely due to various genetic, epigenetic, and demographic mechanisms (Waples 1991; Fraser 2008; Le Luyer et al. 2017; Rodriguez Barreto et al. 2019) that arise a result of adaptation/acclimation to the captive environment (Araki et al. 2007; Christie et al. 2012; Milot et al. 2013; Fraser et al. 2019).

Another major pressure in salmonines, in particular Atlantic salmon, is the escape of farmed fish. Farmed salmon are genetically divergent from wild salmon in a range of traits (Bolstad et al. 2017, 2021), owing to intentional artificial selection for commercially important characteristics, relaxed/domestication selection in captivity, founder effects, and genetic drift (Gjedrem et al. 1991; Gjøen and Bentsen 1997). Escapes from marine fish farms or land-based hatchery units are frequent (Naylor et al. 2005; Jensen et al. 2010). Farmed fish and their hybrids can have substantially reduced fitness in the wild (McGinnity et al. 2003; Skaala et al. 2012; Reed et al. 2015), threatening the genetic integrity and viability of wild populations experiencing introgression (Glover et al. 2017) and altering their life histories (Bolstad et al. 2017, 2021).

Despite the widespread occurrence of anthropogenic hybridisation, whether it be from captive releases, farm escapes, or introductions of closely related conspecifics (e.g., Muhlfeld et al. 2009), considerable variation exists across ecological contexts in the extent of introgression and the magnitude of any associated demographic impacts (White et al. 2018; Lehnert et al. 2020). Density- and frequency- dependent processes are likely key here (Glover et al. 2013; Heino et al. 2015). In particular, the twin concepts of hard and soft selection (Wallace 1975; Bell et al. 2021) are highly relevant, yet rarely considered explicitly. Hard selection refers to situations where the absolute fitness of an individual depends on its phenotype with respect to some environmentally determined optimum. Soft selection, in contrast, occurs when the absolute fitness of an individual depends on its phenotype relative to other conspecifics with which it interacts (Bell et al. 2021). To understand soft selection, it is useful to conceive of the environment as containing a limited number of “ecological vacancies” (Reznick 2016). In order to survive or reproduce, an individual must acquire one of these vacancies, with relative rather than absolute trait values determining which individuals “fill” them. A given trait can be under pure hard selection, pure soft selection, or some combination of the two. Hard selection is independent of, whilst soft selection is dependent upon, the density and phenotypic composition of the population (Bell et al. 2021). To illustrate, consider that body size could be under hard selection if absolute body size determines the match between phenotype and external environment (e.g., thermoregulatory ability), and/or soft selection if relative body size determines success in some intraspecific competition (e.g., resource defence) and there are more competing individuals than vacancies.

Here we present an eco-genetic model to explore the eco-evolutionary consequences of acute and chronic intrusion events by foreign/domesticated individuals into a wild population. Though loosely based on a salmonine lifecycle, the model is generally applicable to any taxon that could experience sequential soft and hard selection events, as well as artificial or natural intrusion by genetically divergent immigrants (e.g., forestry contexts). In our model, individuals compete each generation for a limited number of breeding “slots”, with spawning success determined by a single quantitative trait, *Z_SOFT_*, that is subject to soft selection. Following reproduction, the offspring experience an episode of hard selection wherein survival depends on the match between a second quantitative trait, *Z_HARD_*, and an environmentally-determined trait optimum (with locals assumed to be well-adapted and intruders maladapted). A key prediction we test is that the extent of introgression and its demographic consequences depend on the relative competitiveness of locals versus intruders, i.e., how divergent the two forms are for *Z_SOFT_*. One possibility is that intruders are competitively inferior to locals. For example, experimental studies in salmonines have shown hatchery-bred females to be at a competitive disadvantage relative to wild-bred females at acquiring and defending breeding sites, and hatchery-bred males to be less successful in obtaining mates (Fleming and Gross 1993; Neff et al. 2015). Alternatively, intruders could be competitively superior, if they are for example larger. In the case of commercially cultivated Atlantic salmon, farm escapes are often larger as adults than wild fish, yet they seem to be have lower spawning success (Fleming et al. 1996, 2000; Weir et al. 2004). However, the offspring of farm-farm or farm-wild matings can competitively displace wild-wild offspring, owing to their faster growth rates and hence larger fry sizes (McGinnity et al. 1997, 2003; Fleming et al. 2000). Whilst previous modelling studies have considered genetic and demographic interactions between cultured and wild salmon (Hindar et al. 2006; Baskett et al. 2013; Castellani et al. 2015, 2018; Sylvester et al. 2019; Bradbury et al. 2020), ours is the first, to our knowledge, to explicitly distinguish between hard and soft selection and to explore their interactive effects in this context.

## Methods

### Model description

The model is based on a generalised anadromous salmonine life cycle but, for computational efficiency, with implicit freshwater and saltwater life history stages. The life history is also greatly simplified, to focus directly on the processes of interest (eco-evolutionary interactions between soft and hard selection), without loss of generality. The sequence of model events is as follows: (1) the model is seeded with recruits at the pre-spawner phase; (2) phenotype-dependent competition (soft selection) for limited spawning slots occurs; (3) random mating among spawners and production of new offspring; and (4) offspring survive from the juvenile to the recruit (pre-spawner) stage dependent on the match between phenotype and an environmental optimum (hard selection). Generations are discrete, and time is not explicit within generations.

#### 1. Recruit stage

In all scenarios, the model is seeded with 500 local recruits (*N_R_*(*locals*) = 500) in generation 1, just prior to competition for spawning slots. The initial trait values for *Z_SOFT_* and *Z_HARD_* are a function of the initial allele frequencies *p_SOFT_*(*locals*) and *p_HARD_*(*locals*). Thirty unlinked diploid loci affect each trait (i.e., 60 functional loci in total), and the initial allele frequencies are assumed to be the same across all loci for each trait. In reality, a range of initial allele frequencies could occur (e.g., conforming to a beta distribution; Kardos and Luikart 2021), but this should not affect the qualitative outcomes of the model. The traits are assumed to be initially genetically uncorrelated, although some genetic association between them may emerge over time owing to a build-up of linkage disequilibrium. A third neutral trait is modelled via a single bi-allelic locus, at which locals are assumed to be fixed for a “0” allele and intruders are fixed for a “1” allele. This facilitates the tracking of introgression of neutral foreign alleles into the mixed population over time.

Genotype matrices for each trait for local individuals are established in generation 1. These matrices are 500 rows (individuals) by 60 columns (alleles) in dimension. The first two columns store the alleles for the first locus, the second two columns store the alleles for the second locus, etc. Each element (allele) of the genotype matrix for *Z_SOFT_* was initiated in generation 1 by drawing a number between 0 and 1 from a random uniform distribution, and setting that allele to 1 if the number was less than *p_SOFT_*(*locals*) and 0 if the number was greater than *p_SOFT_*(*locals*). The same procedure was repeated for the genotype matrix for *Z_HARD_*, such that the expected initial allele frequency at each locus equalled *p_HARD_*(*locals*). The genotype matrix for the neutral trait for local individuals was of dimension 500 rows (individuals) by 2 columns (alleles). Local individuals all had a genotype of {0,0} at this neutral diagnostic locus.

In all simulations, the first 20 generations corresponded to a “burn-in” period during which no intrusion occurred. In the acute intrusion scenarios, a given number of intruders (*N_R_*(*intruders*)) was introduced in generation 21 at the recruit stage, with no further intrusion occurring thereafter, whilst in the chronic intrusion scenarios, *N_R_*(*nonlocals*) were introduced in each generation starting from generation 21. The genotype matrix for non-local intruders for *Z_SOFT_* was of dimension *N_R_*(*intruders*) rows by 60 columns (30 diploid loci). The cells of this matrix were populated by drawing a number between 0 and 1 from a random uniform distribution, and setting that allele to 1 if the number was less than *p_SOFT_*(*intruders*) and 0 if the number was greater than *p_SOFT_*(*intruders*). The same process was repeated for the genotype matrix for *Z_HARD_*, such that the expected initial allele frequency for intruders at each locus was *p_HARD_*(*intruders*). Intruders all had a genotype of {1,1} at the neutral diagnostic locus. Immediately after intrusion occurred, the genotype matrices for each trait for locals and intruders were merged by row, such that the new matrices were of dimensions *N_R_*(*total*) rows by 60 columns for *Z_SOFT_* and *Z_HARD_*, and dimension *N_R_*(*total*) rows by two columns for the neutral diagnostic locus, where *N_R_*(*total*) = *N_R_*(*locals*) + *N_R_*(*intruders*).

The genotypic value of each individual for each trait (*Z_SOFT_* and *Z_HARD_*) was then computed by summing the alleles across all 30 loci, assuming that “1” alleles increase the trait value by 1 unit (i.e., the additive allelic effect *α* = 1 at all loci) and “0” alleles have no effect on the trait. Thus, genotypic values ranged from a minimum of 0 to 60. The expected mean genotypic value is equal to 2*npα*, where *n* is the number of loci affecting the trait and *p* is the relevant allele frequency, and the expected genotypic variance is given by 2*np*(1 − *p*)*α*^2^. For example, with *n* = 30, *α* = 1, and *p* = 0.5, the expected mean is 30 and the expected variance is 15. The genotypic means and variances for each trait thus differed between locals and intruders to the extent that *p* differed between them. Non-additive genetic effects were ignored for simplicity, so the genotypic variances corresponded to additive genetic variances (*V*_*A*_).

The initial heritability, ℎ^2^, assumed to be the same for *Z_SOFT_* and *Z_HARD_*, determined the magnitude of the environmental variance (*V*_*E*_) for each trait. *V*_*E*_ was assumed to be constant across generations and was computed as 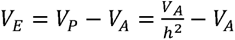, where *V_P_* and *V_A_* were the initial phenotypic and additive genetic variances, respectively. These parameters were defined in generation 1 for local individuals, and in the generation of intrusion for intruders. Note that the actual heritability in any given generation can deviate from initial ℎ^2^, because although *V*_*E*_was assumed to be constant, *V*_*A*_can change under the influence of selection, drift, and introgression. For simplicity, no mutation was included in the model.

#### 2. Soft selection filter

Parental phenotypes are formed for *Z_SOFT_* by drawing an environmental deviation for each individual from a normal distribution of mean 0 and variance equal to *V*_*E*_(*SOFT*), where locals and intruders had potentially different values for the latter parameter, depending on the scenario. Note that in the acute intrusion scenarios, all fish were assumed to be locals from generation 22 onwards, i.e., intrusion of foreign individuals occurred in generation 21 and then any hybrid offspring in future generations were, by definition, locally born. Environmental deviations were added to the genotypic values of individuals, to give individual phenotypic values for *Z_SOFT_*.

The total number of available spawning slots was fixed at *K* = 500 in all simulations. In situations where *N_R_*(*total*) ≤ *K*, all recruits get a spawning slot and no soft selection occurs. In situations where *N_R_*(*total*) > *K* (i.e., when there is reproductive excess), only *K* individuals become spawners, with the surplus assumed to die. To determine which individuals get to spawn, individuals are ranked from top to bottom based on *Z_SOFT_* and only the top *K*/*N_R_*(*total*) fraction of individuals are assigned a spawning slot, which imposes truncational soft selection. For example, if *K* = 500 and *N_R_*(*total*) = 600, only 5 out of every 6 recruits get to spawn, with the top 83% 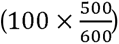 of individuals based on ranked *Z_SOFT_* trait values getting a spawning slot, and the lower 17% failing to spawn. Thus, the higher the reproductive excess (i.e., the more recruits there are relative to spawning slots), the stronger the strength of directional soft selection.

#### 3. Mating and reproduction

Individuals assigned spawning sites then undergo random mating. Separate sexes are not considered, so random hermaphroditic mating based on a classic Wright-Fisher model is assumed. Each individual has an equal chance of becoming a parent, and each individual can produce more than one offspring (or no offspring), which guarantees an approximately Poisson distribution of offspring number per parent (Waples 2022). The total number of new offspring equals *N*_*S*_*k*, where *N*_*S*_ is the number of spawners and *k* the fecundity (number of offspring per parent). In reality, salmonine fishes can produce dozens to thousands of eggs, depending on female size, but for computational efficiency we set *k* = 2. This is effectively equivalent to assuming random mortality of zygotes up to the smolt stage, such that each parent produces an average of two smolts. In other words, all freshwater mortality is subsumed into *k*.

During reproduction, new empty genotype matrices for *Z_SOFT_* and *Z_HARD_*, each of dimension *N*_*O*_ rows by 60 columns, are set up to store the offspring genotypes for each quantitative trait, where *N*_*O*_ = *N*_*S*_*k* is the total number of offspring produced. Similarly, an empty genotype matrix for the neutral trait of dimension *N*_*O*_ rows by two columns is set up. For each offspring, two parents are drawn at random from the pool of *N*_*S*_ spawners by sampling with replacement. For each locus for each trait, the first offspring allele is drawn at random from the two alleles carried by parent 1 at that locus, and the second offspring allele is drawn at random from the two alleles carried by parent 2. This, thus, simulates random segregation and random assortment of alleles into gametes (assuming that loci are unlinked), followed by random fertilisation. This process is repeated across all loci until the new offspring genotype matrices have been fully populated with 1s and 0s.

#### 4. Hard selection filter

In this next step, offspring phenotypes are first formed for *Z_HARD_* by drawing an environmental deviation for each individual from a normal distribution of mean 0 and variance *V*_*E*_(*hard*). As all new offspring are by definition locally-born, regardless of the provenance of their parents, the environmental variance was computed as 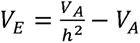, where *V_A_* refers to the initial additive genetic variance of locals in generation 1. Environmental deviations are then added to the genotypic values, to give individual phenotypic values for *Z_HARD_*.

Hard selection is implicitly assumed to occur during the marine phase of the life cycle (although space and life stages are not explicit in the model), notionally corresponding to a scenario where the match between some phenotype (e.g., basal metabolic rate, gape size, growth rate) and marine environment determines marine survival. The expected survival *W*_*i*_ of each individual is computed as a function of its phenotype for *Z_HARD_* based on a Gaussian fitness function:

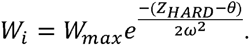

Here, *W*_*max*_ is the maximum survival for individuals whose phenotype coincides with the optimum *θ*, and *ω* corresponds to the “width” of the fitness function in units of phenotypic standard deviations (*sd*(*Z_HARD_*)). In all simulations, *ω* is fixed at a value of 3 × *sd*(*Z_HARD_*)_*t*_, which corresponds to moderate to strong stabilising selection (Estes and Arnold 2007). The *t* subscript here indicates that *sd*(*Z_HARD_*) can vary across generations within model runs, in line with changes in the genetic variance in response to drift, selection and introgression. The alternative was to fix *ω* at a given value across all generations independently of *sd*(*Z_HARD_*)_*t*_, but that would then mean that the effective strength of hard selection would vary as *sd*(*Z_HARD_*) changes over time. A constant strength of selection (for a given deviation of *Z̅*_*HARD*_ from the optimum) was deemed a more parsimonious assumption.

Realised marine survival is then determined by drawing a number between 0 and 1 from a random uniform distribution, with the individual surviving if *W*_*i*_ exceeded this number and dying if the number was less than *W*_*i*_. This imposes hard directional selection on *Z_HARD_* whenever the mean trait value deviates either side of the optimum *θ*. The survivors of hard selection then become the new locally- bred recruits for the next generation, and the model cycles back to step 1.

Each generation, a series of output variables calculated at the recruit stage is stored in a results matrix. These include the phenotypic means and variances for *Z_SOFT_* and *Z_HARD_*, the additive genetic variance for each, the frequency of the foreign ‘1’ allele at the neutral locus, the number of recruits *N_R_*, and the number of spawners *N*_*S*_. The realised recruits per spawner for each generation is then computed as *RPS* = *N_R_*/*N*_*S*_. For all scenarios, 1000 replicate simulations are run and the mean and 95% confidence interval of each output variable of interest is calculated across replicates.

The model was coded in R version 4.3.1 (R Core Team 2023) using the RStudio programming environment (Posit team 2023). All model code is available via GitHub (insert GitHub upon publication).

### Baseline scenarios – no intrusion

To illustrate the basic functionality and behaviour of the model, a series of baseline scenarios was first explored in which no intrusion of foreign fish occurred and the evolutionary and population dynamics were tracked across 100 generations. Initial ℎ^2^for both traits was 0.25, such that each had the potential to respond to selection, and the initial allele frequency at each locus for each trait was 0.75 (i.e., *p_SOFT_* = *p_HARD_* = 0.75). The initial mean values of both traits were rescaled to reference values of 0, corresponding to mean-centering in generation 1. In the first set of baseline simulations, the closed wild population was assumed to be well-adapted with respect to the hard-selected trait, i.e., *Z̅*_*HARD*_ = *θ*. In the second set of baseline simulations, initial maladaptation was assumed (*Z̅*_*HARD*_ < *θ*), such that the hard-selected trait experienced positive directional selection. For simplicity, the optimum *θ* was not allowed to vary over time in each case.

#### Baseline simulations set 1

To illustrate how the strength of soft selection depends on the magnitude of reproductive excess, the ratio of number of recruits *N_R_* to number of spawning slots *K* was varied between ∼1 and ∼1.4. This was achieved by adjusting the *W*_*MAX*_ parameter. Three values of *W*_*MAX*_ were explored: 0.53, 0.63 and 0.73, corresponding to an expected *RPS* of approximately 1.0, 1.2 and 1.4, respectively. When *RPS* ≈ 1 (*W*_*MAX*_ = 0.53), every recruit gained a spawning slot, so there was no reproductive excess. No evolution of *Z_SOFT_* was then expected. When *RPS* ≈ 1.2 (*W*_*MAX*_ = 0.63; moderate reproductive excess), there were more fish than spawning slots, hence phenotype-dependent competition will occur and *Z_SOFT_* should evolve upwards. When *RPS* ≈ 1.4 (*W*_*MAX*_ = 0.73; high reproductive excess), the competition intensified further, and the rate of evolution of *Z_SOFT_* should be correspondingly faster.

#### Baseline simulations set 2

In the second set of baseline simulations, the closed wild population was assumed to be initially poorly adapted to the external environment, by setting *θ* = 20 and keeping initial *Z̅*_*HARD*_ at its reference value of 0. Such a situation could occur if the population experienced a sudden step change in the selective environment, or if it were introduced to a new environment to which it is not pre-adapted (Gomulkiewicz and Holt 1995; Kardos and Luikart 2021). *W*_*MAX*_ was fixed at 0.63, such that expected *RPS* would be greater than 1 in the absence of maladaptation (*θ* = 0) but less than 1 in the presence of maladaptation (*θ* = 20). Thus, the population should initially decline in this scenario but gradually recover towards *K* as *Z_HARD_* evolves towards the new optimum, provided extinction does not occur in the interim, i.e., evolutionary rescue. During the period when *RPS* < 1, no soft selection should occur as all individuals get a spawning slot, but as *RPS* recovers to > 1, soft selection is once again manifested.

### Acute intrusion scenarios

#### Acute intrusion simulations set 1

Here, we assumed that the wild population was initially well-adapted (*Z̅*_*HARD*_(*locals*) = *θ*) and remained closed to intrusion for the first 20 generations, with *W*_*MAX*_ = 0.58 and ℎ^2^(*initial*) = 0.25. Thus, there was some reproductive excess, with ∼550 recruits competing for 500 spawning slots each generation and *RPS* ≈ 1.1. At generation 21, an acute intrusion event occurred wherein 500 foreign/domesticated fish intruded just prior to spawning. The total number of fish competing for spawning slots thus became ∼1050, and soft selection intensified accordingly. From generation 21 onwards, all fish were “locals” in the sense of being locally bred, but many would be of mixed ancestry. The intruders were assumed to be maladapted to the local environmental conditions, such that *Z̅*_*HARD*_(*intruders*) < *Z̅*_*HARD*_(*locals*). This was achieved by setting *p_HARD_*(*intruders*) = 0.25 and *p_HARD_*(*locals*) = 0.75, such that *Z̅*_*HARD*_(*intruders*) was 30 units less than *Z̅*_*HARD*_(*locals*), corresponding to a difference of approximately 4.5 phenotypic standard deviations.

Three scenarios were explored: (1) intruders are competitively inferior to locals (*Z̅*_*SOFT*_ (*intruders*) < *Z̅*_*SOFT*_(*locals*)); (2) intruders are competitively equal to locals (*Z̅*_*SOFT*_(*intruders*) = *Z̅*_*SOFT*_(*locals*)); and (3) intruders are competitively superior to locals (*Z̅*_*SOFT*_ (*intruders*) > *Z̅*_*SOFT*_(*locals*)). This was achieved by varying *p_SOFT_* for intruders relative to locals: in scenario 1, *p_SOFT_*(*intruders*) = 0.4 and *p_SOFT_*(*locals*) = 0.6; in scenario 2, *p_SOFT_*(*intruders*) = *p_SOFT_*(*locals*) = 0.5; and in scenario 3, *p_SOFT_*(*intruders*) = 0.6 and *p_SOFT_*(*locals*) = 0.4. With this parameterisation, *Z̅*_*SOFT*_(*intruders*) was 12 units lower than *Z̅*_*SOFT*_(*locals*) at the time of intrusion in scenario 1, and 12 units higher in scenario 3. This corresponded to an absolute difference in trait means of ∼1.6 phenotypic standard deviations.

#### Acute intrusion simulations set 2

The above simulations were then repeated under a broader range of parameter values, to explore the sensitivity of the results to the level of intrusion and the level of reproductive excess. Three levels of intrusion were explored: low (250 intruders introduced at generation 21); moderate (500 intruders); and high (750 intruders), corresponding to 0.5 × *K*, 1.0 × *K*, and 1.5 × *K*, respectively. Three levels of reproductive excess were also explored: low (*W*_*MAX*_ = 0.53); moderate (*W*_*MAX*_ = 0.58) and high (*W*_*MAX*_ = 0.63), corresponding to expected *RPS* absent any intrusion of ∼1.0, ∼1.1 and ∼1.2, respectively. As before, three levels of relative competitiveness of intruders versus locals were explored (intruders competitively inferior, equal, or superior to locals) using the same parameterisation as the

#### Acute intrusion simulations set 1

This gave a total of 27 scenarios, i.e., combinations of intrusion level, reproductive excess level, and relative competitiveness.

To explore the effects of the level of maladaptation of intruders relative to locals, three additional scenarios were then run in which the difference between *Z̅*_*HARD*_(*intruders*) and *Z̅*_*HARD*_(*locals*) was assumed to be: (1) small (*p_HARD_*(*intruders*) = 0.35; *p_HARD_*(*locals*) = 0.65); (2) moderate (*p_HARD_*(*intruders*) = 0.25; *p_HARD_*(*locals*) = 0.75); and (3) large (*p_HARD_*(*intruders*) = 0.15; *p_HARD_*(*locals*) = 0.85). In all three cases, a moderate level of acute intrusion (500 intruders introduced at generation 21) and a moderate level of reproductive excess (*W*_*MAX*_ = 0.58) was assumed.

### Chronic intrusion scenarios

#### Chronic intrusion simulations set 1

In the chronic intrusion scenarios, a constant number of foreign/domesticated fish were assumed to intrude each generation (from generation 21 onwards) just prior to spawning. In the first set of simulations, the per-generation intrusion rate was fixed at 5% of *K*, where *K*=500. Thus 25 foreign/domesticated fish intruded each generation. As with the acute intrusion simulations, intruders were assumed to be maladapted with respect to *Z_HARD_*, by setting *p_HARD_*(*intruders*) = 0.25 and *p_HARD_*(*locals*) = 0.75. Thus, *Z̅*_*HARD*_ for the intruders is 30 units (∼4.5 phenotypic standard deviations) lower than *Z̅*_*HARD*_ for the locals. As before, the same three levels of relative competitiveness were explored. The initial ℎ^2^ of both traits was set to 0.25 in all cases. The simulations were run for 150 generations, with intrusion starting at generation 21.

#### Chronic intrusion simulations set 2

In the second set of chronic intrusion simulations, all parameters were the same as in set 1, except the per-generation intrusion rate was increased to 20% of *K*.

#### Chronic intrusion simulations set 3

In the final set of simulations, a broader range of parameter values was explored. Specifically, the sensitivity of the chronic intrusion results to trait heritability and the level of reproductive excess was tested. In these simulations, the per-generation intrusion rate was set to 10% of *K*. Two levels of trait heritability (same value applies to both *Z_SOFT_* and *Z_HARD_*) were explored: ℎ^2^(*initial*) = 0.25 and ℎ^2^(*initial*) = 0.50. Three levels of reproductive excess were also explored: low (*W*_*MAX*_ = 0.53); moderate (*W*_*MAX*_ = 0.58) and high (*W*_*MAX*_ = 0.63), corresponding to an expected *RPS* absent any intrusion of circa 1.0, 1.1 and 1.2, respectively. Again, the same three levels of relative competitiveness of intruders versus locals were explored. This gave a total of 18 scenarios, i.e., combinations of trait heritability, reproductive excess level, and relative competitiveness.

## Results

### Baseline scenarios

#### Baseline simulations set 1

The results of the first set of baseline simulations showed that, as expected, the rate of evolution of *Z_SOFT_* depended on the extent of reproductive excess (Fig.1). With low reproductive excess, little to no directional evolution of *Z_SOFT_* occurred (Fig.1A, orange curve) because all, or nearly all, recruits gained a spawning slot each generation, with *RPS* fluctuating around 1. Because soft selection only occurs when there are more recruits than spawning slots, this creates an asymmetric situation where a small amount of soft selection will occur whenever *RPS* is by chance >1 (*N_R_* > *K*), but not when it is by chance <1 (*N_R_* < *K*). This meant that a small amount of evolution of *Z_SOFT_* accrued across multiple generations in the low reproductive excess case, which explains why the orange curve in Fig.1A shifted slightly upwards over time. With moderate reproductive excess (green curve in Fig.1A), the rate of evolution of *Z_SOFT_* was faster, and with high reproductive excess (pink curve in Fig.1A) it was faster again. In all three scenarios, no evolution of *Z_HARD_* occurred (Fig.1B) because the population was well adapted (*Z̅*_*HARD*_ coincided with *θ*).

**Fig. 1.**
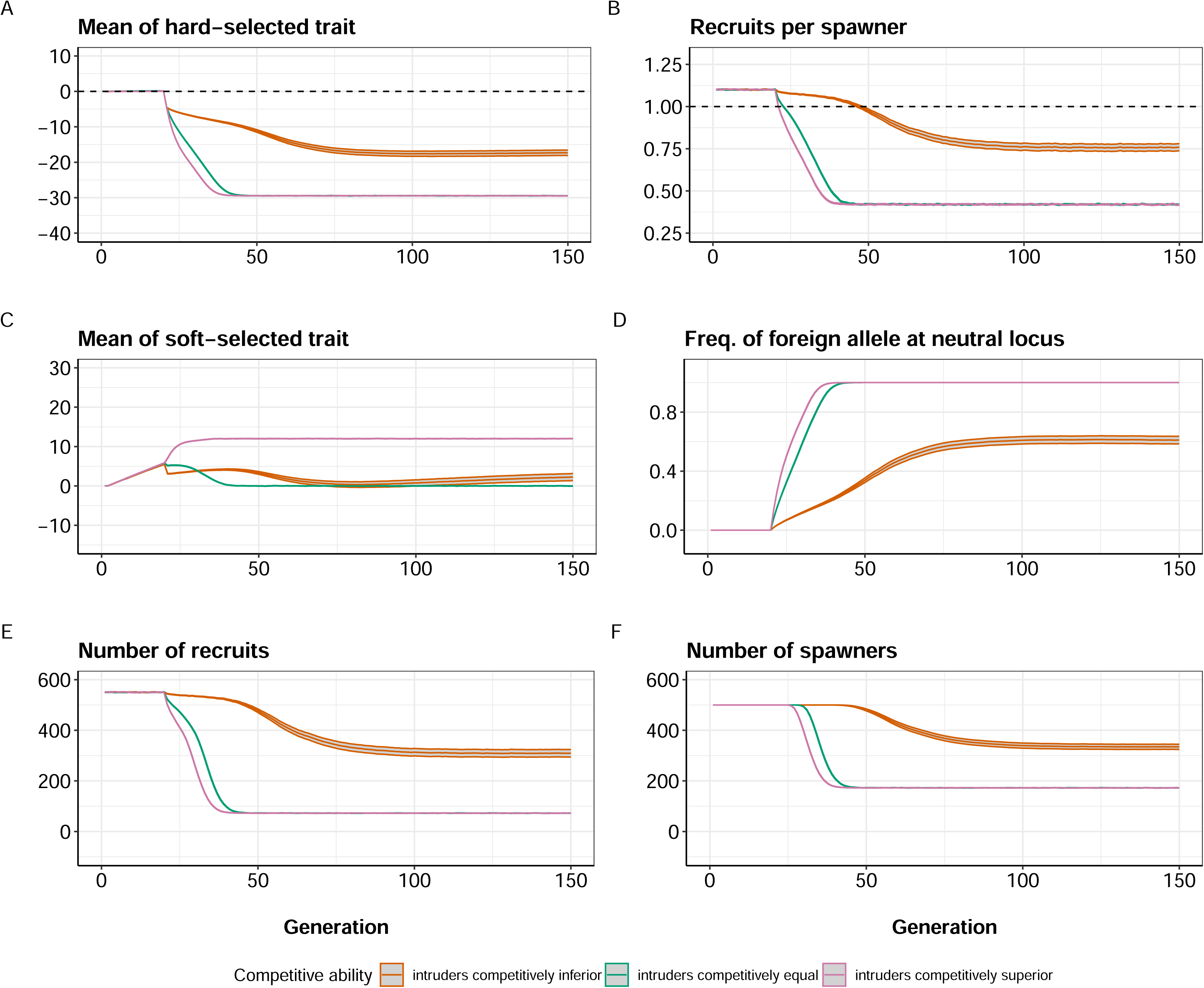
Results of baseline simulations set 1. No intrusion of foreign/domesticated fish occurred, and the population was assumed to be initially well-adapted with respect to the hard-selected trait. Orange = low reproductive excess (*W*_*MAX*_ = 0.53); green = moderate reproductive excess (*W*_*MAX*_ = 0.63); pink = high reproductive excess (*W*_*MAX*_ = 0.73). Mean and 95% confidence intervals (grey ribbons) across 1000 replicate simulations shown. (A) Changes in mean of soft-selected trait (*Z̅*_*SOFT*_) over time. (B) Changes in mean of hard-selected trait (*Z̅*_*HARD*_) over time (dashed line indicates optimum). (C) Changes in genetic variance of *Z_SOFT_* over time. (D) Changes in genetic variance of *Z_HARD_* over time.

With moderate or high reproductive excess, the rate of evolution of *Z_SOFT_* gradually plateaued, as selection limits were reached due to the erosion of genetic variation. With only 30 loci contributing to each trait, allelic variation will be lost even under pure drift (in the absence of mutation or gene flow), as loci go to fixation for a given allele. Thus, additive genetic variation in *Z_SOFT_* went down slowly over time in the scenario with low reproductive excess (largely drift only) and more rapidly in the moderate and high reproductive excess scenarios (selection + drift; Fig.1C). *Z_HARD_* experienced no directional selection, so the rate of loss of genetic variance (Fig.1D) in all three scenarios was similar to that for *Z_SOFT_* in the low reproductive excess scenario.

#### Baseline simulations set 2

Here, the wild population was again closed to intrusion but now the optimum *θ* was 20 units higher than *Z̅*_*HARD*_ in generation 1. As a result of this maladaptation, *Z_HARD_* evolved upwards towards *θ* (Fig.2A). The rate of evolution of *Z_HARD_* slowed down as the generations progressed for two reasons: (1) selection weakened as the new optimum was approached (because the Gaussian fitness landscape is flatter near the optimum), and (2) genetic variance was progressively lost owing to both directional selection and drift (Fig.S1).

**Fig. 2:**
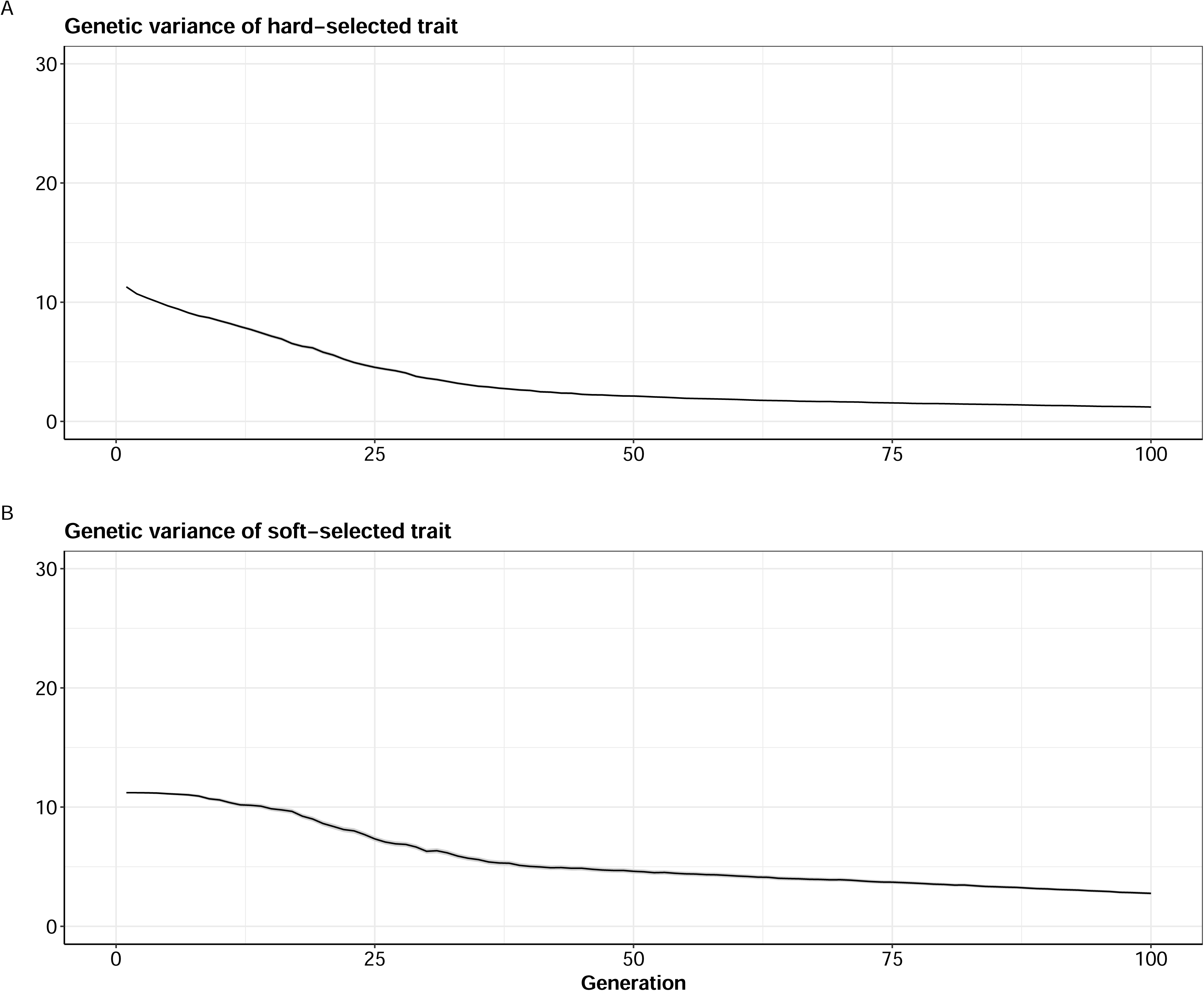
Results of baseline simulations set 2. No intrusion of foreign/domesticated fish occurred, and the population was assumed to be initially maladapted with respect to the trait optimum (initial *Z̅*_*HARD*_ = 0; *θ* = 20). Moderate reproductive excess (*W*_*MAX*_ = 0.63) was assumed and initial ℎ^2^ = 0.25. Mean and 95% confidence intervals (grey ribbons) across 1000 replicate simulations shown. (A) Evolutionary trajectory of *Z_HARD_* (dashed line = optimum). (B) Changes in recruits per spawner (*RPS*) over time where the dashed line represents exact population replacement (i.e., no increase/decrease between generations). (C) Evolutionary trajectory of *Z_SOFT_*. (D) Trajectory of number of spawners *N*_*S*_over time. In all panels, averages taken each generation over only those replicate populations that persisted (*N*_*S*_ > 0).

Up until around generation 25, *RPS* < 1 owing to the maladaptation (Fig.2B), so no soft selection occurred and correspondingly no directional evolution of *Z_SOFT_* was observed (Fig.2C). During this early period of maladaptation, the number of spawners declined but then recovered in a classic U-shape pattern under evolutionary rescue (Fig.2D) as positive population growth was restored. Twenty percent of replicate populations went extinct in the interim. From about generation 25, *RPS* had recovered to above 1 (Fig.1C), and soft selection began acting upon *Z_SOFT_*, which gradually evolved upwards thereafter (Fig.2D).

### Acute intrusion scenarios

#### Acute intrusion simulations set 1

Prior to intrusion, *Z_SOFT_* evolved gradually upwards from generation 1 to 20 (Fig.3A), as there were approximately 1.1 recruits for every spawner (i.e., reproductive excess). *Z_HARD_* remained static during this pre-intrusion phase (Fig.3B), as the optimum was constant. The subsequent effects of acute intrusion depended strongly on the relative competitiveness of intruders and locals. When intruders were competitively equal to locals (green curves in Fig.3), consistent soft selection occurred and so *Z_SOFT_* evolved gradually upwards (Fig.3A, green curve). When intruders were competitively inferior to locals, *Z_SOFT_* exhibited a sudden drop in generation 21 when intrusion occurred (Fig.3A, orange curve). However, *Z_SOFT_* rapidly jumped back up in the ensuing few generations, because any mixed ancestry individuals with lower *Z_SOFT_* would have experienced a strong selective disadvantage (reduced likelihood of attaining a spawning site). *Z_SOFT_* continued to evolve gradually upwards thereafter, as ongoing competition and soft selection played out.

**Fig. 3:**
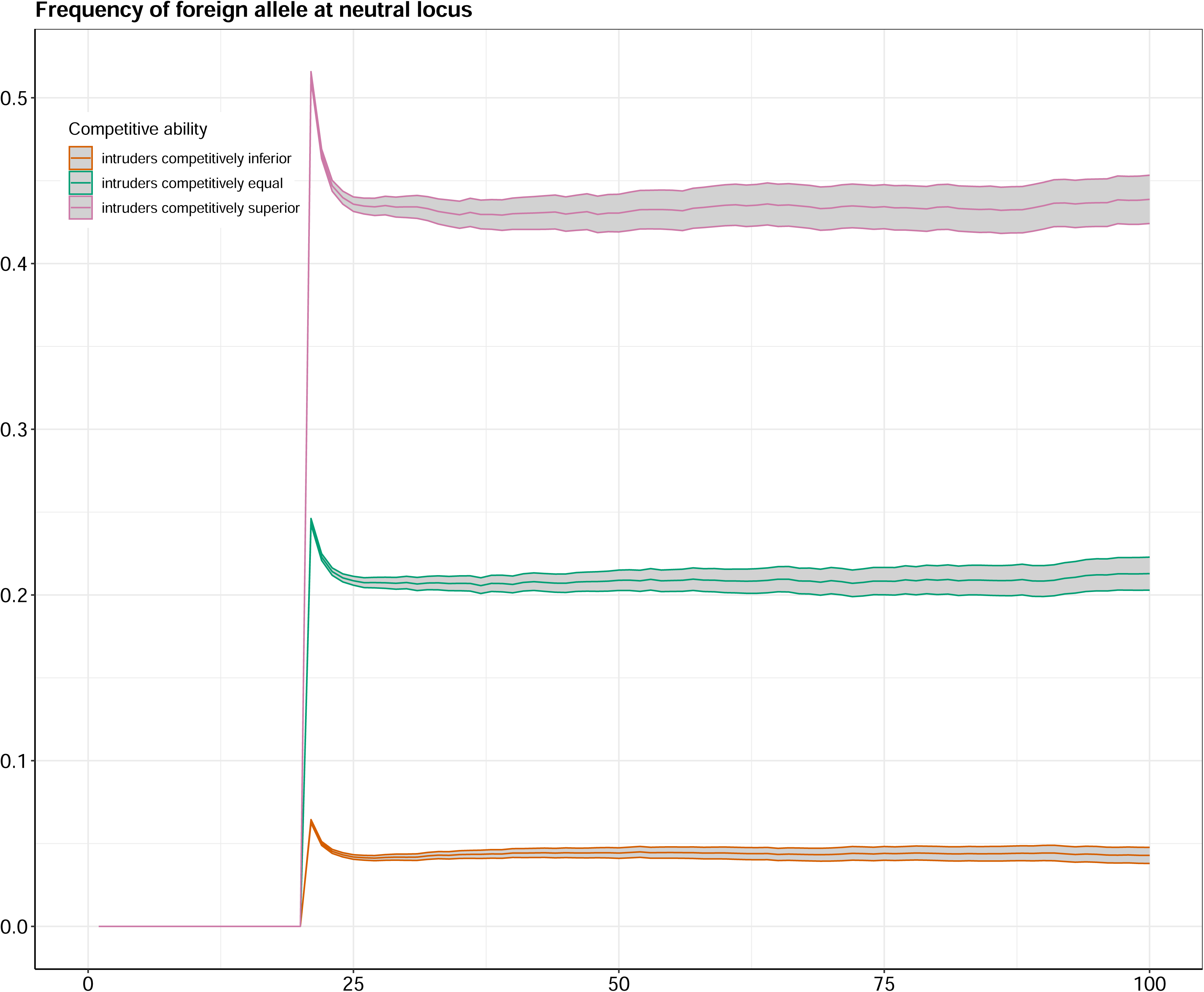
Results of acute intrusion simulations set 1. Prior to intrusion in generation 20, the wild population has a *RPS* of ∼1.1 (*W*_*MAX*_ = 0.58), such that ∼550 recruits compete for *K*= 500 spawning slots. At generation 20, 500 foreign/domesticated fish intrude just prior to spawning, giving ∼1050 fish in total, greatly intensifying competition for the 500 spawning slots. Results of three scenarios (mean and 95% confidence intervals across 1000 replicates) are shown: orange = intruders competitively inferior to locals; green = intruders competitively equal to locals, and pink = intruders competitively superior to locals. The intruders are maladapted with respect to *Z_HARD_* in all three cases, so *RPS* goes down in generation 20 but then slowly recovers as *Z̅*_*HARD*_ of the mixed population evolves back up towards the fixed optimum (*θ* = 0; dashed line in B). (A) Evolutionary trajectory of *Z_SOFT_*. (B) Evolutionary trajectory of *Z_HARD_*. (C) Trajectory of *RPS* over time (dashed line = replacement). (D) Trajectory of number of spawners *N*_*S*_ over time. Initial ℎ^2^= 0.25.

In contrast, when intruders were competitively superior, *Z_SOFT_* rapidly jumped up in generation 21 as a direct result of the intrusion (Fig.3A, pink curve). For the next 25 generations or so, *Z_SOFT_* remained relatively static, because little soft selection occurred. The latter reflected the fact that *Z_HARD_* was pulled strongly off its optimum by introgression (Fig.3B, pink curve). The effect of this maladaptation is seen in the much lower dip below 1 that was exhibited by *RPS* compared to the other two scenarios (Fig.3C, pink curve). The greater maladaptation in this scenario was due to the highly competitive intruders (and their hybrid/backcrossed descendants in the generations immediately post-intrusion) having high spawning success, which led to higher introgression of foreign/domesticated alleles into the population (Fig.S2). The net result was that the number of spawners in this scenario dipped to a much lower nadir compared to the scenarios where intruders were competitively equal or inferior to locals (Fig.3D). Nevertheless, evolutionary rescue occurred in all three scenarios.

#### Acute intrusion simulations set 2

The basic patterns found in the first set of acute intrusion simulations were emulated in the second set, with the effects scaling with the degree of intrusion and the degree of reproductive excess. The higher the intrusion rate, the greater the negative impact of acute intrusion on the number of spawners (Fig.4, compare bottom panels to top panels). The number of spawners was reduced to a lower level when the intrusion rate was higher, because *Z_HARD_* was dragged more from the optimum (Fig.S3). In contrast, the greater the reproductive excess, the weaker the negative impact of acute intrusion on population size (Fig.4, compare right panels to left panels), because *N_R_* remained above *K* for longer (Fig.S4). The level of reproductive excess did not affect the level of maladaptation (Fig.S3), but rather the effects of a given level of maladaptation on the number of spawners. *RPS* was negatively affected by intrusion in all cases, but this translated into strong impacts on number of spawners only in those scenarios where *RPS* was reduced below 1 (Fig.S4). For a given level of intrusion and reproductive excess, intrusion had a greater negative impact on the number of spawners when intruders were competitively superior. The probability of extinction (fraction of replicate populations that went extinct) was higher when the level of intrusion was higher, when the level of reproductive excess was lower, and when intruders were competitively superior to locals (Fig.S5).

**Fig. 4:**
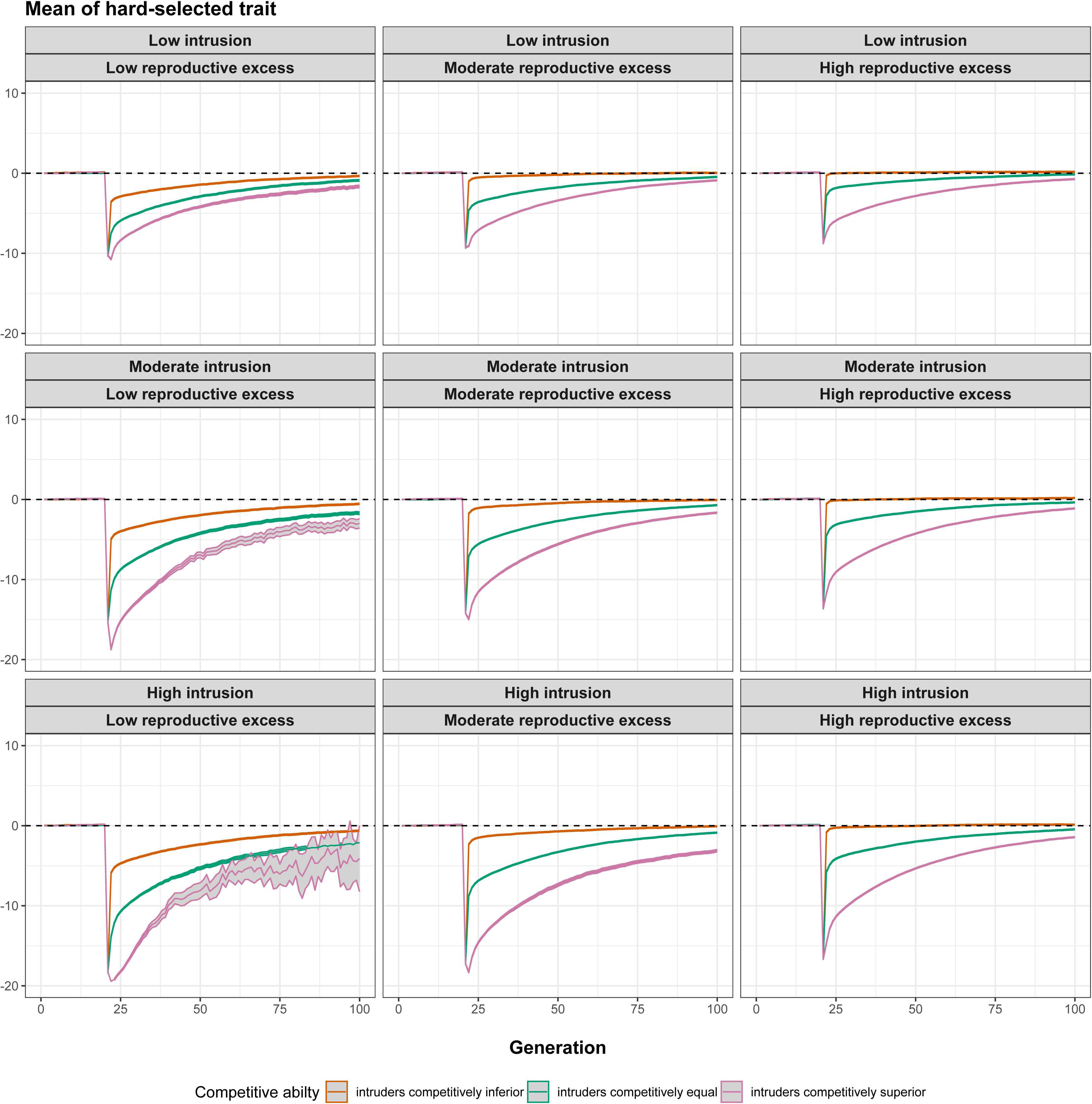
Results of acute intrusion simulations set 2. Mean and 95% confidence intervals across 1000 replicates shown. Orange = intruders competitively inferior to locals; green = intruders competitively equal to locals, and pink = intruders competitively superior to locals. *K*=500 in all scenarios. Low intrusion = 250 intruders introduced in generation 21; moderate intrusion = 500 intruders introduced; high intrusion = 750 intruders introduced. Low reproductive excess: *W*_*MAX*_ = 0.53; moderate reproductive excess: *W*_*MAX*_ = 0.58; high reproductive excess: *W*_*MAX*_ = 0.63. Each panel shows the trajectory of number of spawners over time, with the average taken each generation over only those replicate populations that persisted (*N*_*S*_ > 0). Initial ℎ^2^= 0.25.

The patterns were the same (results not shown) when *Z̅*_*HARD*_(*intruders*) was instead assumed to be greater, rather than less than, *Z̅*_*HARD*_(*locals*), because the adaptive landscape was symmetrical about the optimum. The patterns were also more pronounced when a bigger absolute difference between *Z̅*_*HARD*_(*intruders*) and *Z̅*_*HARD*_(*locals*) was assumed (i.e., greater maladaptation of intruders), and less pronounced when a smaller absolute difference was assumed (i.e., weaker maladaptation of intruders; Fig.S6).

### Chronic intrusion scenarios

#### Chronic intrusion simulations set 1

Chronic intrusion exerted a consistent downwards pull on *Z_HARD_*, because the intruders were maladapted to the local environmental conditions. This was counteracted, however, by an upwards pull, as evolution “tried” to bring *Z_HARD_* back towards the fixed optimum (*θ* = 0). When the intruders were competitively equal to the locals, *Z_HARD_* gradually evolved downwards towards a value of -30 (Fig.5A, green curve), because chronic intrusion resulted in the effective “genetic extinction” of the wild population. As the degree of maladaptation went up, *RPS* went down accordingly, approaching a minimum around 0.5 (Fig.5B, green curve). *Z_SOFT_* went up initially, during the pre-intrusion period (generations 1 to 20), as positive soft selection was occurring. Once *RPS* dipped below 1 by around generation 25, however, soft selection no longer occurred and hence *Z_SOFT_* was dragged downwards (Fig.5C, green curve) by the continual influx of foreign/domesticated fish each generation. Once *Z_SOFT_* approached the reference value of 0, it remained there as soft selection was no longer occurring given that *RPS* < 1. The frequency of the foreign/domesticated allele at the neutral locus increased steadily in this scenario and reached fixation by around generation 100 (Fig.5D, green curve), indicating effective genetic replacement of the original wild population by the intruders. The number of recruits declined to <50 by around generation 100 (Fig.5E), with the number of spawners levelling out at around 50-70 (Fig.5F).

**Fig. 5:**
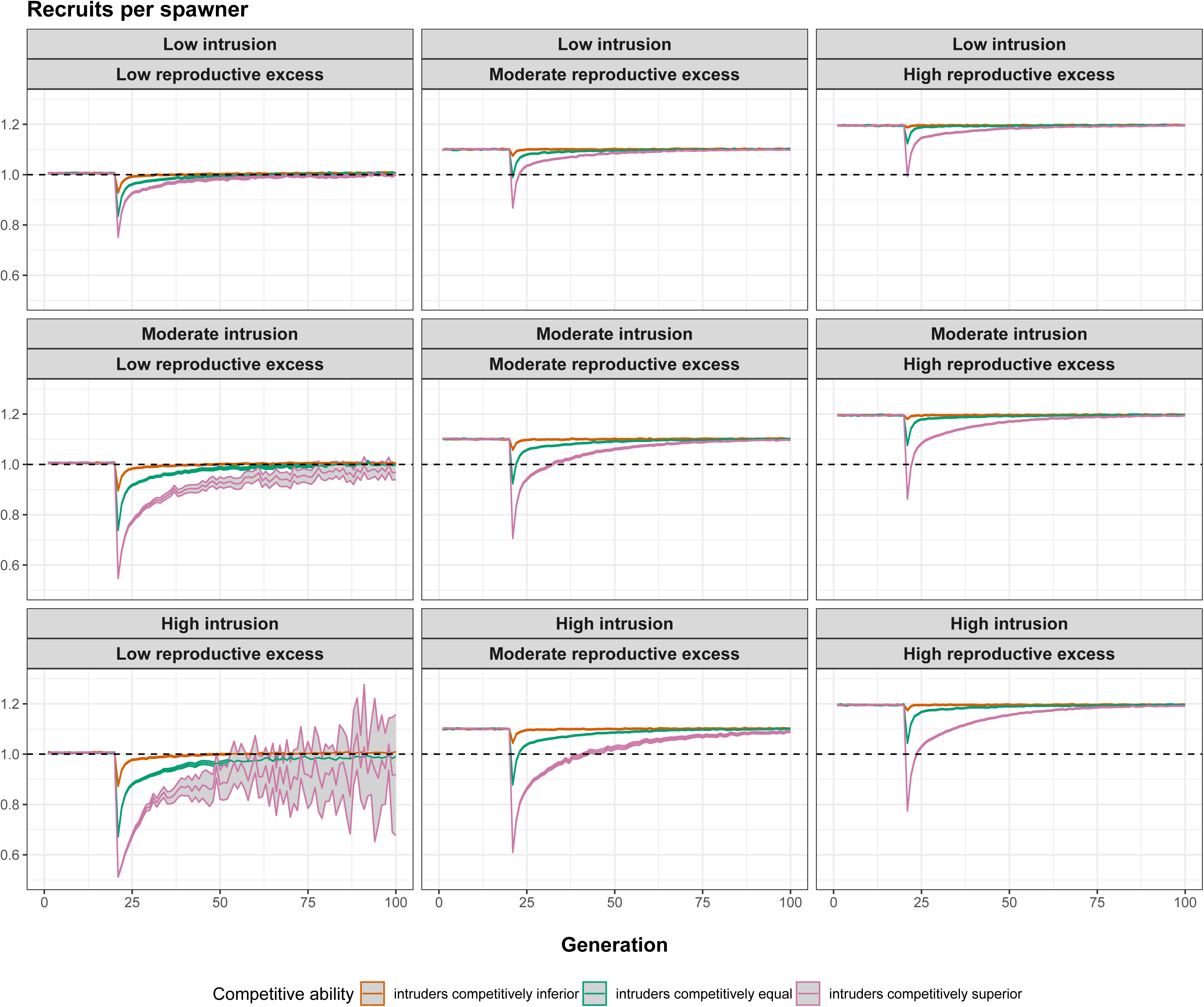
Results of chronic intrusion simulations set 1. Prior to intrusion in generation 21, the wild population has a *RPS* of ∼1.1 (*W*_*MAX*_ = 0.58), such that ∼550 recruits compete for *K*= 500 spawning slots. From generation 21 onwards, 25 foreign/domesticated fish intrude each generation just prior to spawning. Results of three scenarios (mean and 95% confidence intervals across 1000 replicates) are shown: orange = intruders competitively inferior to locals; green = intruders competitively equal to locals, and pink = intruders competitively superior to locals. Each panel shows the trajectory of number of spawners over time, with the average taken each generation over only those replicate populations that persisted (*N*_*S*_ > 0). Initial ℎ^2^= 0.25.

When intruders were competitively superior, the results were similar to the “intruders competitively equal” scenario, except that maladaptation increased faster (Fig.5A, pink curve) and hence *RPS* declined faster (Fig.5B, pink curve). *Z_SOFT_* stabilized around +10 (Fig.5C, pink curve), reflecting the fact that gene swamping again occurred so that the original wild population was replaced genetically by foreign/domesticated alleles (Fig.5D, pink curve). Only around 20-25 recruits were produced each generation (Fig.5E, pink curve), and the number of spawners per generation bottomed out at around 30- 40 (Fig.5F, pink curve).

The results were very different when intruders were competitively inferior to locals. Soft selection filtered out most intruders in each generation, such that little maladaptation occurred (Fig.5A, orange curve). As a result, *RPS* remained steady above 1 (Fig.5B, orange curve). *Z_SOFT_* continued to evolve upwards (Fig.5C, orange curve). Very little introgression of foreign/domesticated alleles occurred (Fig.5D, orange curve), and the number of recruits remained steady at around 550 (Fig.5E, orange curve) and the number of spawners remained at *K* = 500 (Fig.5F, orange curve).

#### Chronic intrusion simulations set 2

The results of the second set of chronic intrusion simulations (in which 100 foreign/domesticated fish intruded each generation) were similar to the first set (in which only 25 intruded), except *Z_HARD_* declined faster towards its equilibrium value (Fig.6A). In the ‘intruders competitively equal’ and ’intruders competitively superior’ scenarios, complete genetic replacement of locals by the foreign/domesticated type occurred (Fig.6D, green and pink curves), and the number of spawners equilibrated at just under 200 (Fig.6F, green and pink curves). With the higher intrusion rate, some introgression occurred even in the ‘intruders competitively inferior’ scenario (Fig.6D, orange curve), indicative of a hybrid swarm situation. Some maladaptation occurred (Fig.6A, orange curve), albeit less than in the ‘intruders competitively equal’ and ‘intruders competitively superior’ scenarios (Fig.6A, green and pink curves respectively). *RPS* equilibrated at a value below 1 (Fig.6B, red curve). The number of spawners stabilized at around 350 (Fig.6F, orange curve), which was considerably higher than the ‘intruders competitively equal’ and ‘intruders competitively superior’ scenarios (Fig.6F, green and pink curves respectively).

**Fig. 6:**
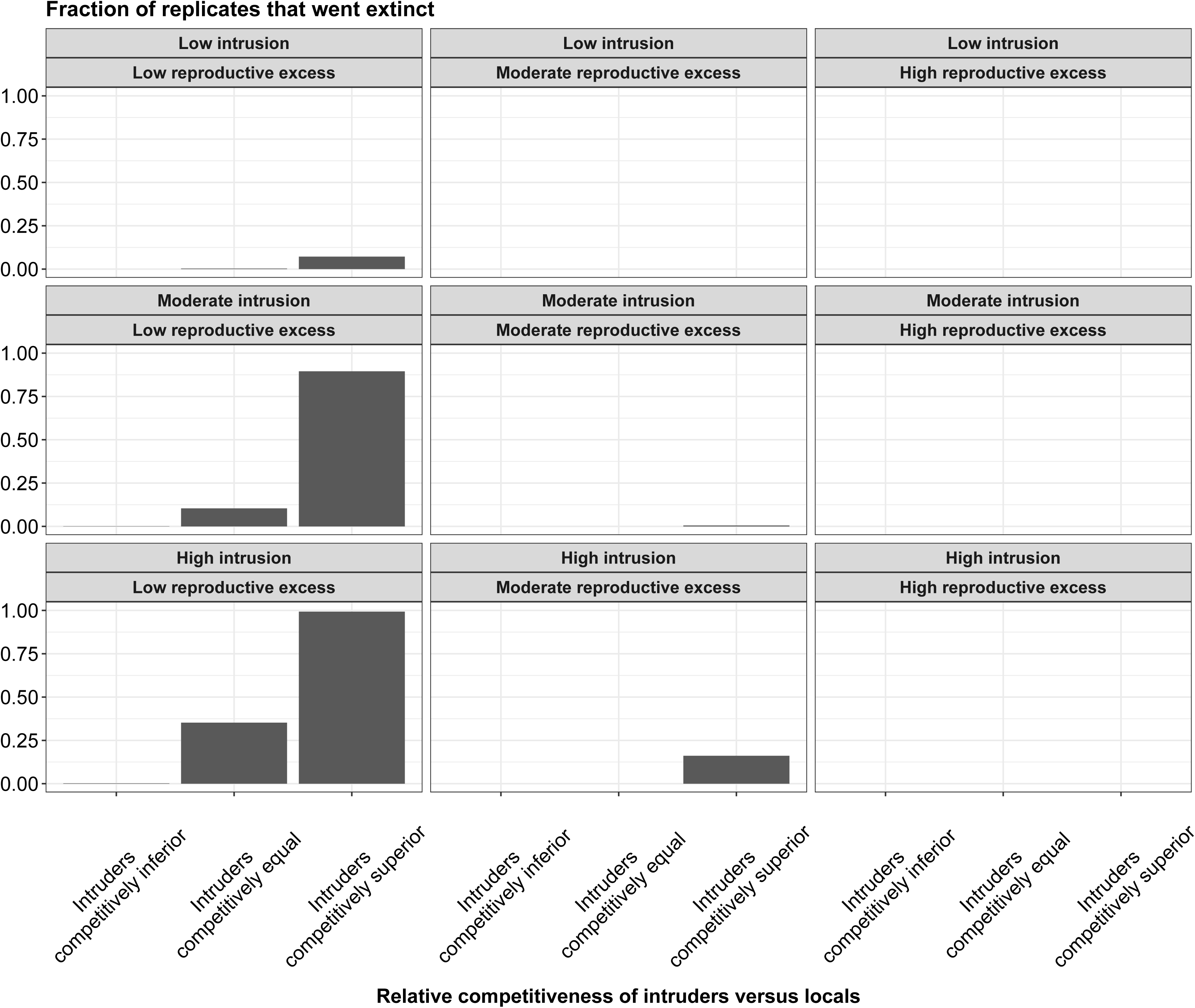
Results of chronic intrusion simulations set 2. Prior to intrusion in generation 20, the wild population has a *RPS* of ∼1.1 (*W*_*MAX*_ = 0.58), such that ∼550 recruits compete for *K*= 500 spawning slots. From generation 20 onwards, 100 foreign/domesticated fish intrude each generation just prior to spawning. Results of three scenarios (mean and 95% confidence intervals across 1000 replicates) are shown: orange = intruders competitively inferior to locals; green = intruders competitively equal to locals, and pink = intruders competitively superior to locals. The intruders are maladapted with respect to *Z_HARD_* in all three cases. (A) Evolutionary trajectory of *Z_HARD_* (dashed line = optimum). (B) Trajectory of *RPS* over time (dashed line = replacement). (C) Evolutionary trajectory of *Z_SOFT_*. (D) Changes in frequency of foreign/domesticated allele at neutral locus over time. (E) Trajectory of number of recruits *N_R_* over time. (F) Trajectory of number of spawners *N*_*S*_ over time. Initial ℎ^2^= 0.25.

#### Chronic intrusion simulations set 3

The results of the chronic intrusion scenarios were sensitive to both the trait heritability and the degree of reproductive excess. In the low reproductive excess scenario (Fig.S7), population size was reduced to 100 or fewer spawners by generation 50 or so. Recruitment at this point was close to zero, so new spawners each generation effectively consisted of fresh waves of intruding immigrants. Trait heritability had little effect on these outcomes.

The dynamics changed when there was moderate reproductive excess (Fig.S8). When trait heritability was 0.25, the results were similar to the low reproductive excess scenario, but only in the ‘intruders competitively equal’ and ‘intruders competitively superior’ scenarios. In contrast, under the ‘intruders competitively inferior’ scenario, soft selection filtered out most intruders each generation so little maladaptation or introgression occurred. As a result, population size remained stable at *K*. When heritability was increased to 0.5, the outcomes were the same as the heritability = 0.25 case for the ‘intruders competitively superior’ and ‘intruders competitively inferior’ scenarios, but different for the ‘intruders competitively equal’ scenario (Fig.S8). In the latter case, continued intrusion resulted in a small amount of maladaptation, but not enough to cause an appreciable decrease in *RPS*, so the number of spawners remained close to *K*.

The dynamics changed yet again when there was high reproductive excess (Fig.S9). When trait heritability was 0.25, the results were similar to the moderate reproductive excess scenario, but only in the ‘intruders competitively superior’ scenario. Strong maladaptation occurred, and population size was reduced by about generation 50 to 100 or fewer spawners. In contrast, in the ‘intruders competitively equal’ and especially the ‘intruders competitively inferior’ scenarios, soft selection filtered out many intruders each generation. The number of spawners remained stable at *K* in both cases. When heritability was increased to 0.5, little to no introgression or maladaptation occurred in the ‘intruders competitively inferior’ and ‘intruders competitively equal’ scenarios, whilst a small amount of introgression and maladaptation occurred in the ‘intruders competitively superior’ scenario (Fig.S9), with the number of spawners remaining stable at *K* in all three cases.

## Discussion

Our modelling results showed that the eco-evolutionary consequences of maladaptive hybridisation depend on the relative competitiveness of intruders versus locals. In both the acute and chronic intrusion scenarios, competitive superiority of intruders over locals acted as a ‘Trojan Horse’ for accelerated introgression of foreign alleles into the admixed population, reducing the average degree of adaptation (match between *Z_HARD_* and the optimum) and population productivity (recruits per spawner). Under certain parameterisations, this led to population decline (acute intrusion scenarios) or genetic replacement of local genotypes by non-local genotypes (chronic intrusion scenarios). However, these negative outcomes were much less pronounced, or absent altogether, when intruders were competitively inferior to locals. In that case, soft selection effectively cushioned the wild population against maladaptive hybridisation by filtering out foreign genotypes each generation at the spawning stage and hence limiting the scope for introgression. Taken together, these findings emphasise how complex interactions between hard and soft selection play a crucial role in determining the evolutionary and demographic implications of hybridisation between divergent gene pools.

Soft selection remains a relatively poorly appreciated aspect of eco-evolutionary dynamics, yet one that is highly relevant to a range of applied problems including artificial propagation, farm escapes, climate change, harvest and habitat alterations (Bell et al. 2021). The term soft selection has been mostly used in the context of models of evolution in spatially heterogeneous environments or meta-populations (e.g., Levene 1953; Christiansen 1975; Ravigné et al. 2004; Ho and Agrawal 2012; Hadfield and Reed 2022), in which density regulation is local and the contribution of each habitat/deme to total population size is fixed. Here, however, we did not consider spatial heterogeneity or metapopulation structure; our focus was on a single undivided population. We thus use the term soft selection as originally conceived by Wallace (1975), in that the expected fitness of an individual with a given trait value for *Z_SOFT_* depended on both the number of other conspecifics and their phenotypic composition for this trait. The density-dependent nature of soft selection can be seen clearly in our baseline scenarios, where greater reproductive excess (more recruits relative to spawning slots) was associated with faster evolution of *Z_SOFT_* (Fig. 1A). *Z_HARD_*, in contrast, was unaffected by the magnitude of reproductive excess (Fig.1B), because hard selection is independent of both population density and composition (Bell et al. 2021).

In our simulations, we assumed that *Z_SOFT_* and *Z_HARD_* were genetically independent traits, but in reality they could be genetically correlated owing to pleiotropy or linkage disequilibrium (Walsh and Lynch 2018). Direct selection on one would then cause a correlated (indirect) response in the other, and their evolutionary trajectories would no longer be independent. Pleiotropy could be modelled in future extensions by allowing some fraction of the total number of loci to be shared among the traits (e.g., Castellani et al. 2015; Kane et al. 2022). Even if the traits are independent at a genetic level, however, selection on *Z_HARD_* can still influence the strength of selection on *Z_SOFT_*. For example, if the deviation between *Z̅*_*HARD*_ and the optimum increases owing to environmental change or influx of maladapted genotypes, this will reduce average survivorship and hence population productivity. This, in turn, will weaken soft selection because fewer recruits now compete for spawning slots. Such an effect was apparent in the *Baseline simulations set 2*, where no selection on *Z_SOFT_* occurred for the first 25 generations or so, during which the population was strongly maladapted (following an assumed step change in the environment), because *RPS* was less than 1. Once *RPS* was restored by evolutionary rescue to >1 (via evolution of *Z_HARD_*), soft selection was again observed (Fig. 2C). The outcomes would be more complex if *Z_SOFT_* and *Z_HARD_* were genetically correlated, as soft selection would then have indirect consequences for hard selection, whilst hard selection would have both direct and indirect consequences for soft selection.

Acute intrusion of maladapted invaders had similar consequences to a sudden shift in the optimum caused by environmental change in the absence of intrusion. That is, *Z̅*_*HARD*_ was dragged away from the optimum, such that directional selection on *Z_HARD_*, and potential evolutionary rescue, then ensued. The key difference, however, was that intrusion altered the competitive dynamics whenever *Z_SOFT_* of intruders and locals was different. When the intruders were competitively inferior (*Z̅*_*SOFT*_(*intruders*) < *Z̅*_*SOFT*_(*locals*)), many of them simply failed to spawn, and hence the demographic penalty on the admixed population owing to maladaptation of *Z_HARD_* was much lower, relative to a scenario where intruders and locals were competitively equal (Fig.3). In contrast, competitive superiority of intruders over locals exacerbated maladaptation and increased extinction risk. These effects scaled, in turn, with the degree to which intruders were locally maladapted (differed in mean *Z_HARD_*) relative to locals (Fig.S6). Similar results were found in our chronic intrusion scenarios, except that “genetic extinction” rather than demographic extinction was more likely to occur when the intruders were competitively superior (Figs. 5&6). The productivity (recruits per spawner) of the admixed population was still greatly depressed in the chronic intrusion scenarios, however, relative to cases where intruders were competitively inferior or equal.

Acute or chronic intrusions can occur under both hatchery release and farm escape contexts. For example, stocking of rivers or lakes with hatchery fish (and their potential subsequent straying into neighbouring catchments) might occur on a once-off or intermittent pulse basis, or more regularly on an annual basis. Similarly, escapes from fish farms might involve large episodic events (e.g. Sylvester et al. 2019), or continual low-level “drip leakage” (Glover et al. 2017). Whether, and by how much, the intruders differ from locals in competitive ability will depend on the specific types of traits and life stages involved, and the extent to which soft/density-dependent selective pressures under artificial propagation differ from those in the wild. In our model, we assumed that competition occurred over access to limited spawning sites, which has certainly been a major factor in the historical evolution of salmonine life histories (Young 2004). Hatchery-bred salmonines tend to be competitively inferior to wild-bred fish at acquiring/defending breeding sites and mates, which may reflect a relaxation of natural and sexual selection in captivity (Fleming and Gross 1993; Neff et al. 2015). Similarly, escapees from commercial aquaculture facilities (fish farms) seem to be at a competitive disadvantage (particularly males) during spawning under natural conditions, although their spawning success relative to wild fish varies across contexts depending on, for example, the life stage at which the fish escape (Fleming et al. 1996, 2000; Weir et al. 2004). The specific traits mediating these spawning interactions, and their genetic basis, remain largely unknown, but behavioural phenotypes might be more important than body size, given that farmed adults are often larger than wild adults and hence should be superior competitors on that basis alone. However, competition for limited fry territories might also be important in determining whether farm genes can or cannot introgress into wild populations. Selection in the farm typically favours rapid growth, so the offspring of farm escapes are typically larger as fry than offspring of wild fish, with hybrids intermediate (see Glover et al. 2017 and references therein). As a result, pure farm or hybrid offspring can competitively displace pure wild offspring to poorer quality habitats where survival is lower (McGinnity et al. 1997, 2003; Fleming et al. 2000). Whilst we did not model such a scenario, competitive superiority of intruder genotypes at the fry/parr stage (Sundt-Hansen et al. 2015) would presumably have similar qualitative effects in our model as competitive superiority at the spawning stage, because soft selection and hard selection would still interact in the same way.

Our modelling framework is similar in some ways, but fundamentally different in others, to previous modelling studies that have considered genetic and demographic interactions between cultured and wild fish (Hindar et al. 2006; Baskett et al. 2013; Castellani et al. 2018; Sylvester et al. 2019; Bradbury et al. 2020). Although not an explicitly eco-genetic model (*sensu* Dunlop et al. 2009), the model of Hindar et al. (2006) did explore the effects of varying spawning success of escaped farmed salmon relative to wild salmon. The results suggested that low relative spawning success of farm escapees substantially reduces the proportion of farmed genotypes in the admixed population in subsequent generations, similar to our finding that competitive inferiority of intruders leads to lower introgression. They also showed that spawning by mature male parr may act as a conduit for gene flow from farmed to wild salmon (Garant et al. 2003), an interesting complication that we did not consider in our model but which clearly also involves density and frequency dependent processes (Kane et al. 2022). The eco-genetic model of Baskett et al. (2013) showed that an intermediate degree of maladaptation of aquaculture escapees relative to wild fish has the most serious consequences, because extremely maladapted escapees are purged before they get a chance to reproduce. For a given degree of maladaptation, the mean fitness taken across all individuals in the admixed population was higher, and the recovery rate faster, when density dependence in their model occurred after selection (what they called soft selection) rather than before (what they called hard selection). However, selection itself was not density/frequency dependent in their model (i.e., not soft selection as defined by Wallace 1975). Nor did Baskett et al. (2013) model potential competitive superiority of cultured fish over wild fish, which they argued (in their discussion) would likely “*increase the demographic effect of aquaculture escapees on wild populations and the importance of the relative timing of escape and density dependence*”. In our model, density dependence and (soft) selection occur simultaneously at the spawning stage, but adding a subsequent round of ecological (i.e., phenotype-independent) density dependence either before or after hard selection would likely lead to similar findings as Baskett et al. (2013).

The IBSEM model of Castellani et al. (2015) is much more mechanistic and specifically tailored to salmon biology than our model (see also Reed et al. 2011; Piou and Prévost 2012), and is thus better suited to applications where the goal is to generate predictions specific to a given stock or region (e.g., Castellani et al. 2018; Sylvester et al. 2019; Bradbury et al. 2020). IBSEM incorporates density- dependent growth and survival at different life stages, but interactive effects of genotype and density on these vital rates are not included, to the best of our knowledge. Thus, selection is effectively assumed to be hard. Like Hindar et al. (2006), IBSEM assumes that domesticated escapees have lower spawning success (30% for females, 5% for males) relative to wild spawners, which effectively limits introgression of farm alleles, but competitive abilities (during spawning or any other life stage) are genotype independent. It would be interesting to see how the eco-evolutionary dynamics predicted by IBSEM would be affected by including soft selection dynamics. Likewise, future extensions of our model could consider additional complexities such as mutation, simultaneous gene flow from multiple farm strains (Besnier et al. 2011), dispersal of farm escapes (Bradbury et al. 2020), straying from nearby wild populations (Bradbury et al. 2020), mature male parr (Hindar et al. 2006; Castellani et al. 2015), correlational selection (Tufto 2010), or uneven distribution of allelic effects across loci (Castellani et al. 2015; Kardos and Luikart 2021). However, increasing model complexity can come at the cost of generality and interpretability, so the appropriate amount of biological detail will depend on the specific research questions being addressed.

## Conclusions

We have demonstrated that considering interactions between soft and hard selection can be crucial in evaluating the potential eco-evolutionary consequences of influxes of genetically divergent intruders into wild populations. Our model broadens the scope of previous modelling studies (Hindar et al. 2006; Tufto 2010; Baskett et al. 2013; Castellani et al. 2015, 2018; Sylvester et al. 2019; Bradbury et al. 2020) by allowing for genotype-dependent variation in competitive abilities, which may be a key determinant of introgression levels. Aquaculture escapes, in particular, are recognised as an ongoing threat to the productivity and persistence of wild salmonine populations (Forseth et al. 2017; Glover et al. 2017), and our results inform aquaculture risk assessments and salmon conservation more generally. They also emphasise the importance of obtaining better information across a range of ecological or invasion contexts on the relative competitiveness of domesticated/foreign individuals and their wild counterparts. Climate change will likely exacerbate the consequences of maladaptive hybridisation, so better knowledge of the processes influencing a population’s ability to resist intrusion should ultimately foster evolutionarily-informed conservation decisions.

## Acknowledgements

This work was supported by funding from Science Foundation Ireland, the Marine Institute, and the Department for the Economy, Northern Ireland under the Investigators Programme Grant Number SFI/15/IA/3028. TER was funded by an ERC Starting Grant (639192) and an SFI ERC Support Award. RJOS was supported by a Suomen Akatemia Profi7 award (Human Diversity – Award Number: 352727) and the Investigators Programme Grant Number SFI/15/IA/3028. PMG was supported by the Investigators Programme Grant Number SFI/15/IA/3028 and the Marine Institute, Ireland. The authors declare no conflicts of interest with the work herein.

---

**Fig. S1.**
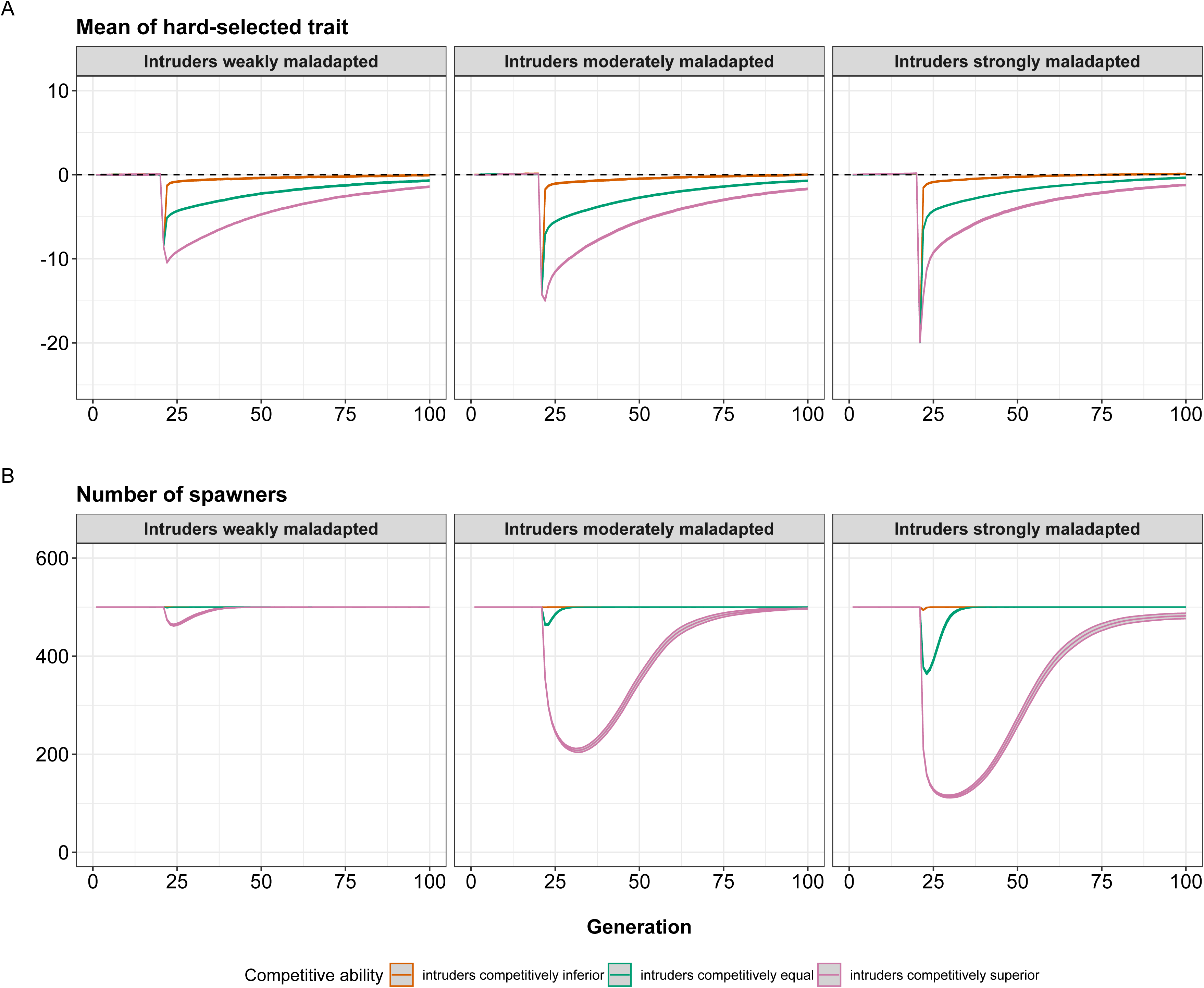
Changes in additive genetic variance in (A) *Z_HARD_* and (B) *Z_SOFT_* over time in the baseline simulations set 2. Mean and 95% confidence intervals (grey ribbons) across 1000 replicate simulations shown.

**Fig. S2:**
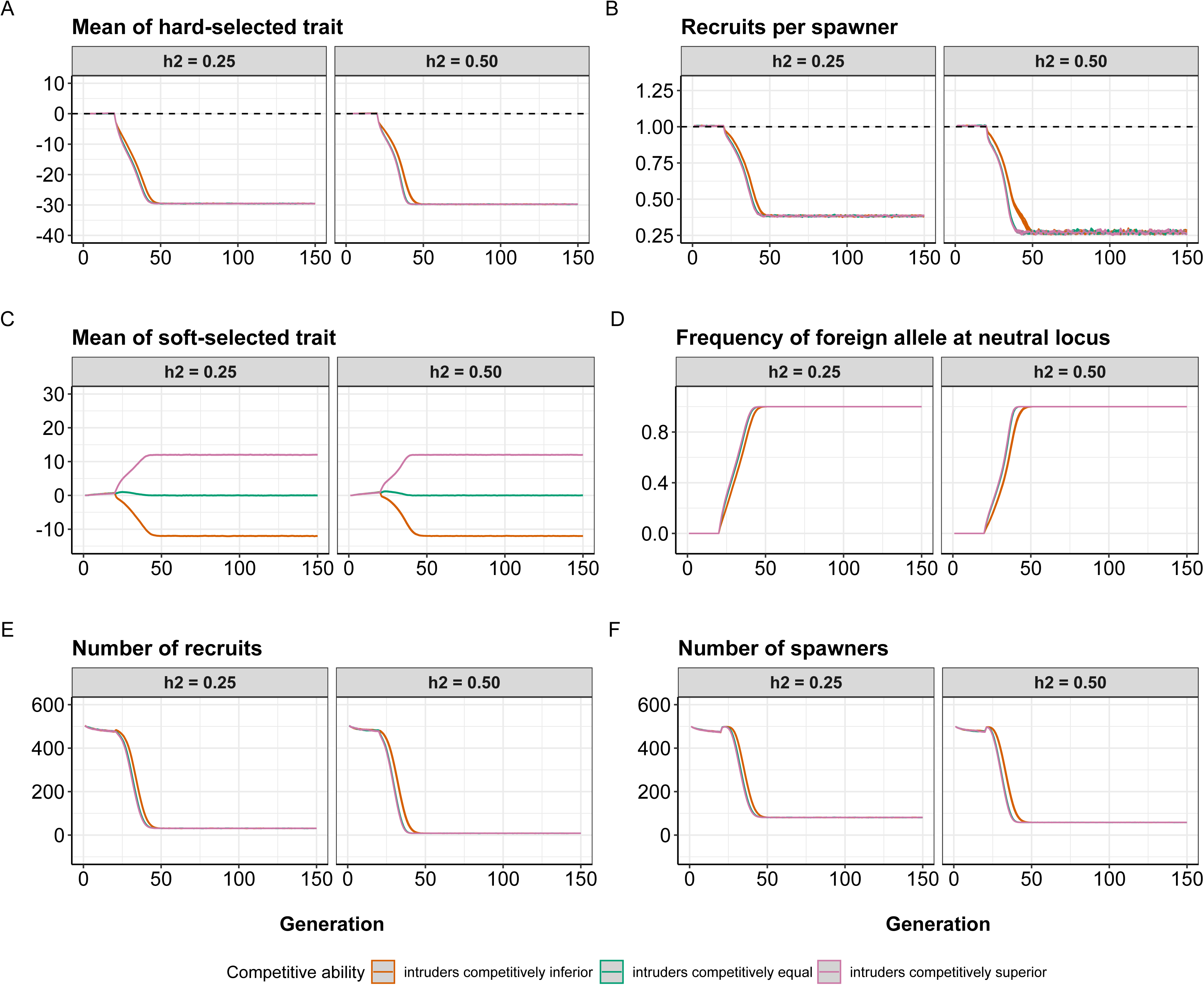
Dynamics of introgression at the neutral locus in the acute intrusion simulations set 1. The y-axis shows the frequency of the foreign/domesticated allele at the single neutral locus. Mean and 95% confidence intervals (grey ribbons) across 1000 replicate simulations shown.

**Fig. S3:**
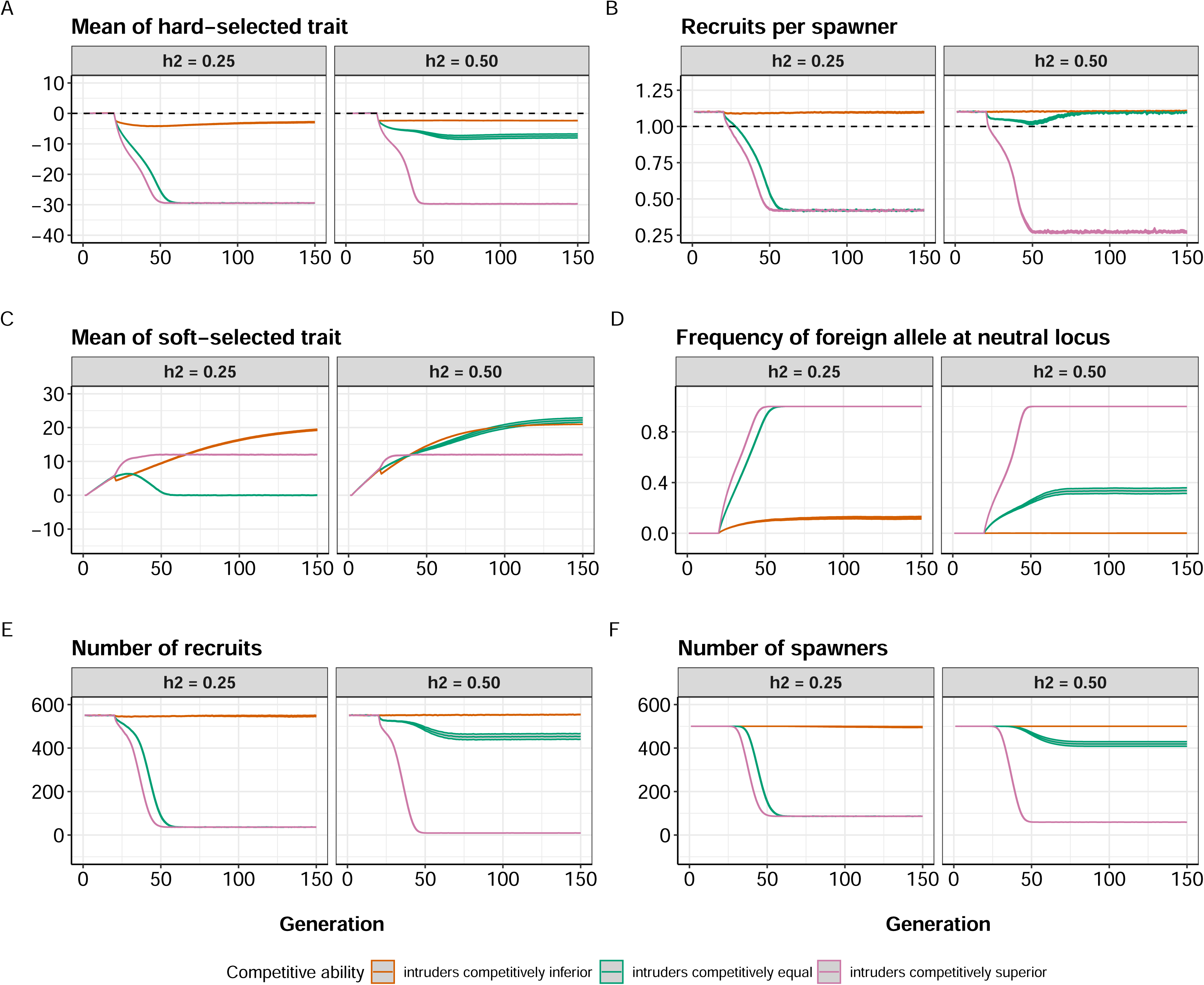
The evolutionary dynamics of *Z_HARD_* in the intrusion simulations set 2. *K*=500 in all scenarios. Low intrusion = 250 intruders introduced in generation 20; moderate intrusion = 500 intruders introduced; high intrusion = 750 intruders introduced.

**Fig. S4:**
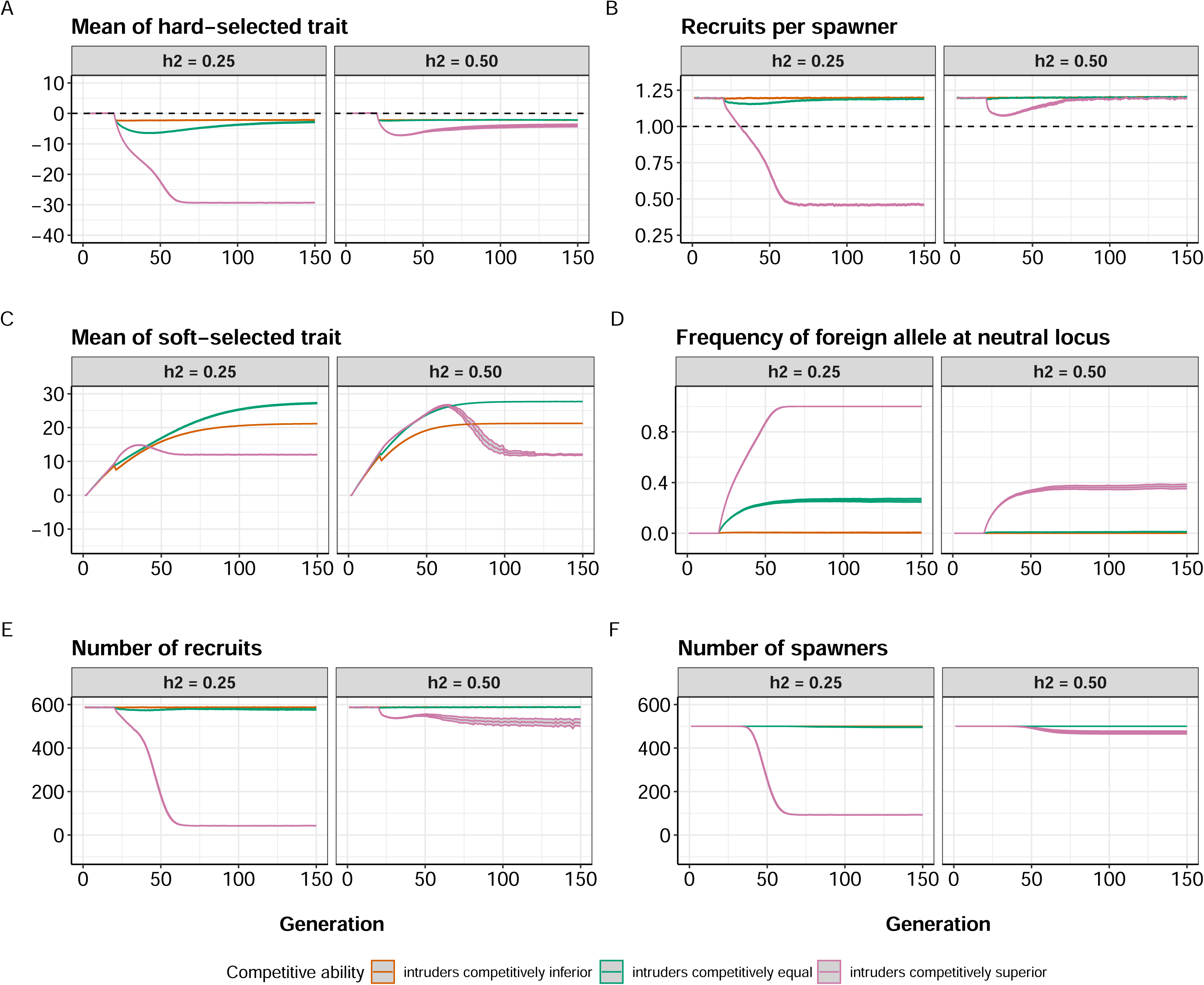
Changes in *RPS* in the acute intrusion simulations set 2. *K*=500 in all scenarios. Low intrusion = 250 intruders introduced in generation 20; moderate intrusion = 500 intruders introduced; high intrusion = 750 intruders introduced. Low reproductive excess: *W*_*MAX*_ = 0.53; moderate reproductive excess: *W*_*MAX*_ = 0.58; high reproductive excess: *W*_*MAX*_ = 0.63. Each panel shows the trajectory of *RPS* over time, with the average taken each generation over only those replicate populations that persisted (*N*_*S*_ > 0). Initial ℎ^2^= 0.25.

**Fig. S5:**
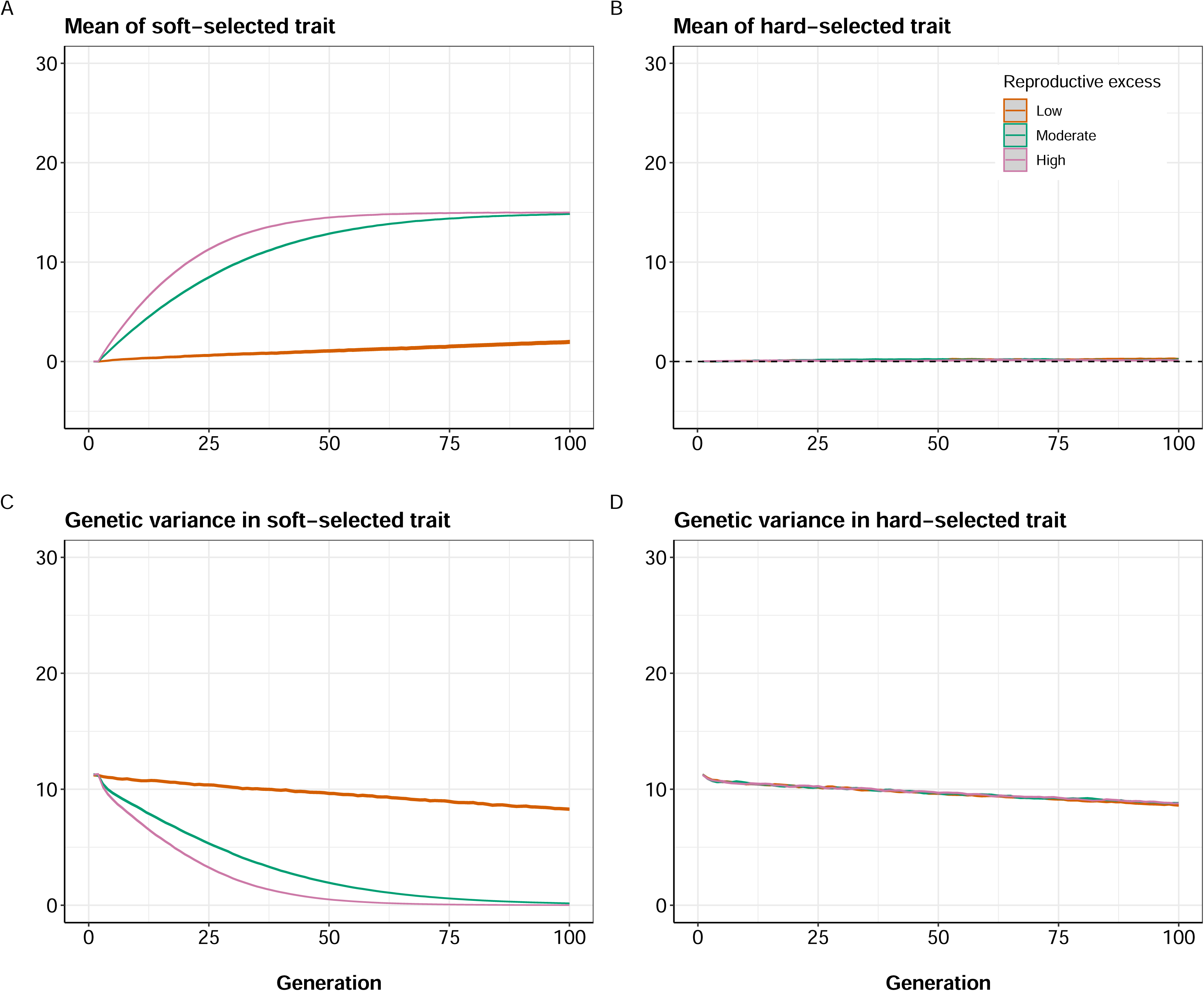
Probability of extinction in the acute intrusion simulations set 2. *K*=500 in all scenarios. Low intrusion = 250 intruders introduced in generation 20; moderate intrusion = 500 intruders introduced; high intrusion = 750 intruders introduced. Low reproductive excess: *W*_*MAX*_ = 0.53; moderate reproductive excess: *W*_*MAX*_ = 0.58; high reproductive excess: *W*_*MAX*_ = 0.63. Initial ℎ^2^= 0.25.

**Fig. S6:**
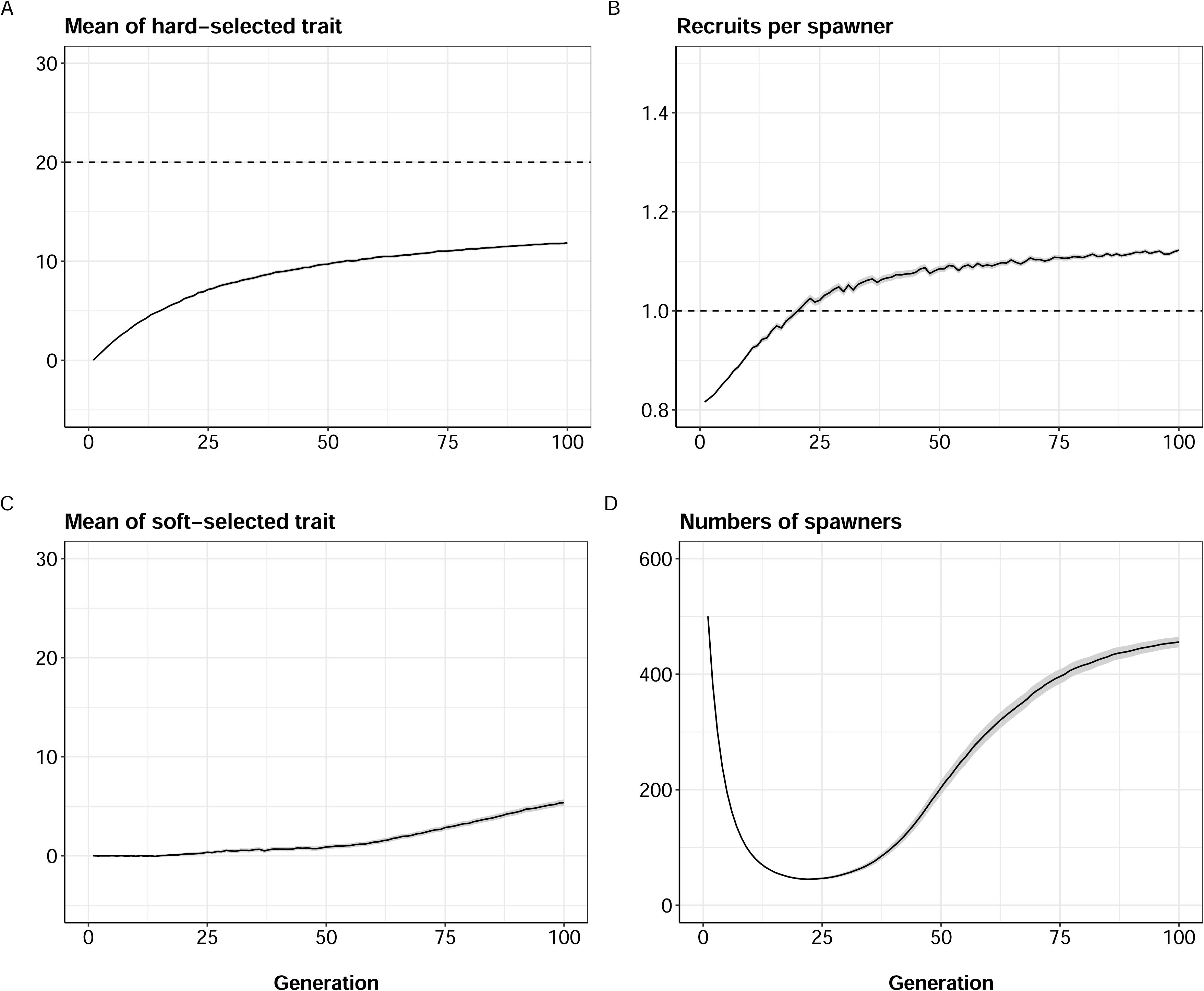
Effects of level of maladaptation of intruders on the results of acute intrusion simulations. (A) Evolutionary trajectory of *Z_HARD_*. (B) Number of spawners through time. Mean and 95% confidence intervals across 1000 replicates shown. In all cases, a moderate level of acute intrusion (500 intruders introduced at generation 20) and a moderate level of reproductive excess (*W*_*MAX*_ = 0.58) was assumed, with initial ℎ^2^= 0.25.

**Fig. S7:**
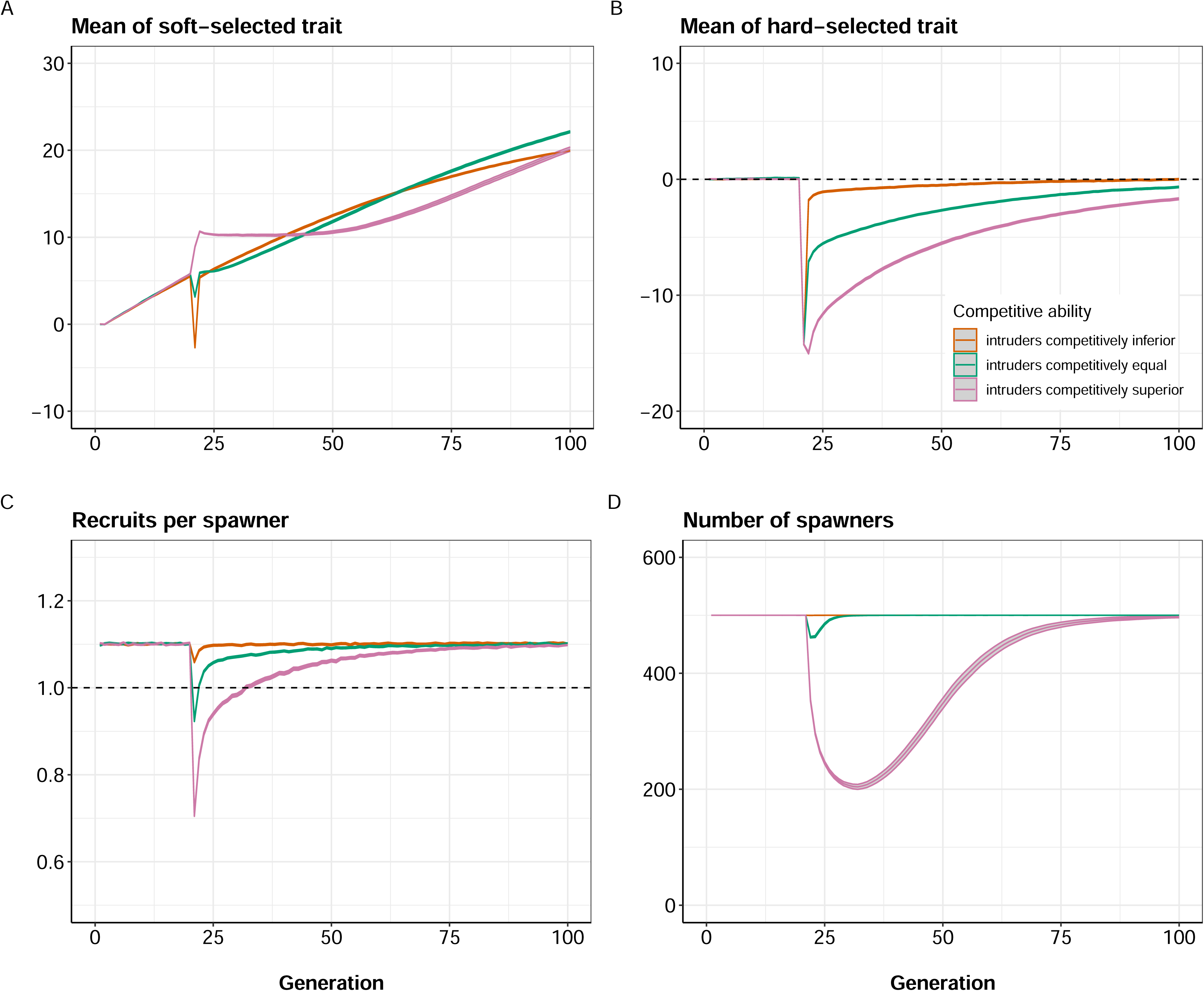
Results of chronic intrusion simulations set 3 for the low reproductive excess scenario (*W*_*MAX*_ = 0.53). Each panel shows the results (mean and 95% confidence intervals across 1000 replicate simulations) comparing cases where the initial heritability of both *Z_SOFT_* and *Z_HARD_* was ℎ^2^ = 0.25 (left sub-panels) or ℎ^2^ = 0.50 (right sub-panels). The per-generation intrusion rate was fixed at 10% of *K*, i.e., 50 foreign/domesticated fish intruded each generation.

**Fig. S8:**
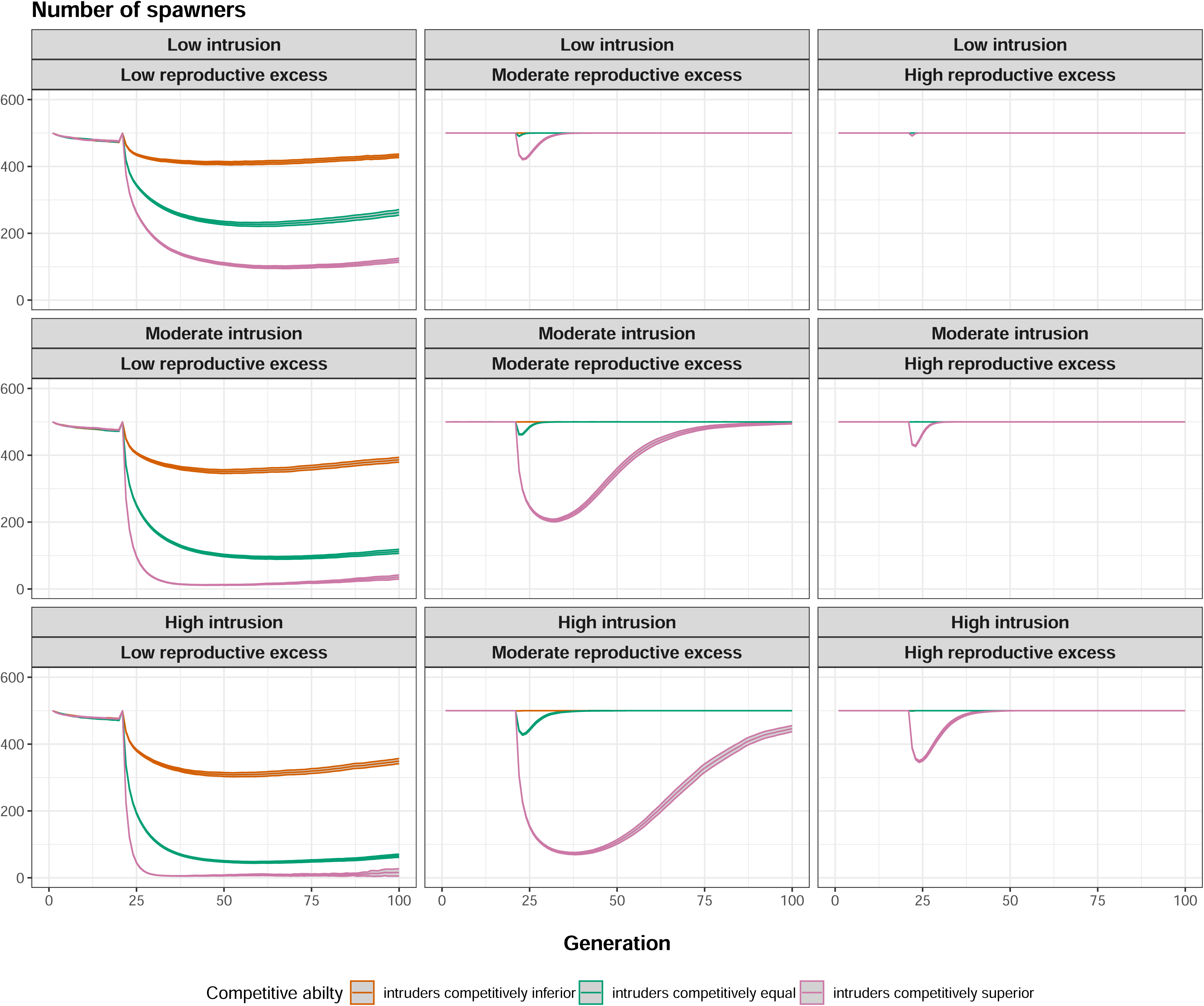
Results of chronic intrusion simulations set 3 for the moderate reproductive excess scenario (*W*_*MAX*_ = 0.58). Each panel shows the results (mean and 95% confidence intervals across 1000 replicate simulations) comparing cases where the heritability (*h^2^*) of both *Z_SOFT_* and *Z_HARD_* = 0.25 (left sub-panels) or= 0.50 (right sub-panels). The per-generation intrusion rate was fixed at 10% of *K*, i.e., 50 foreign/domesticated fish intruded each generation.

**Fig. S9:**
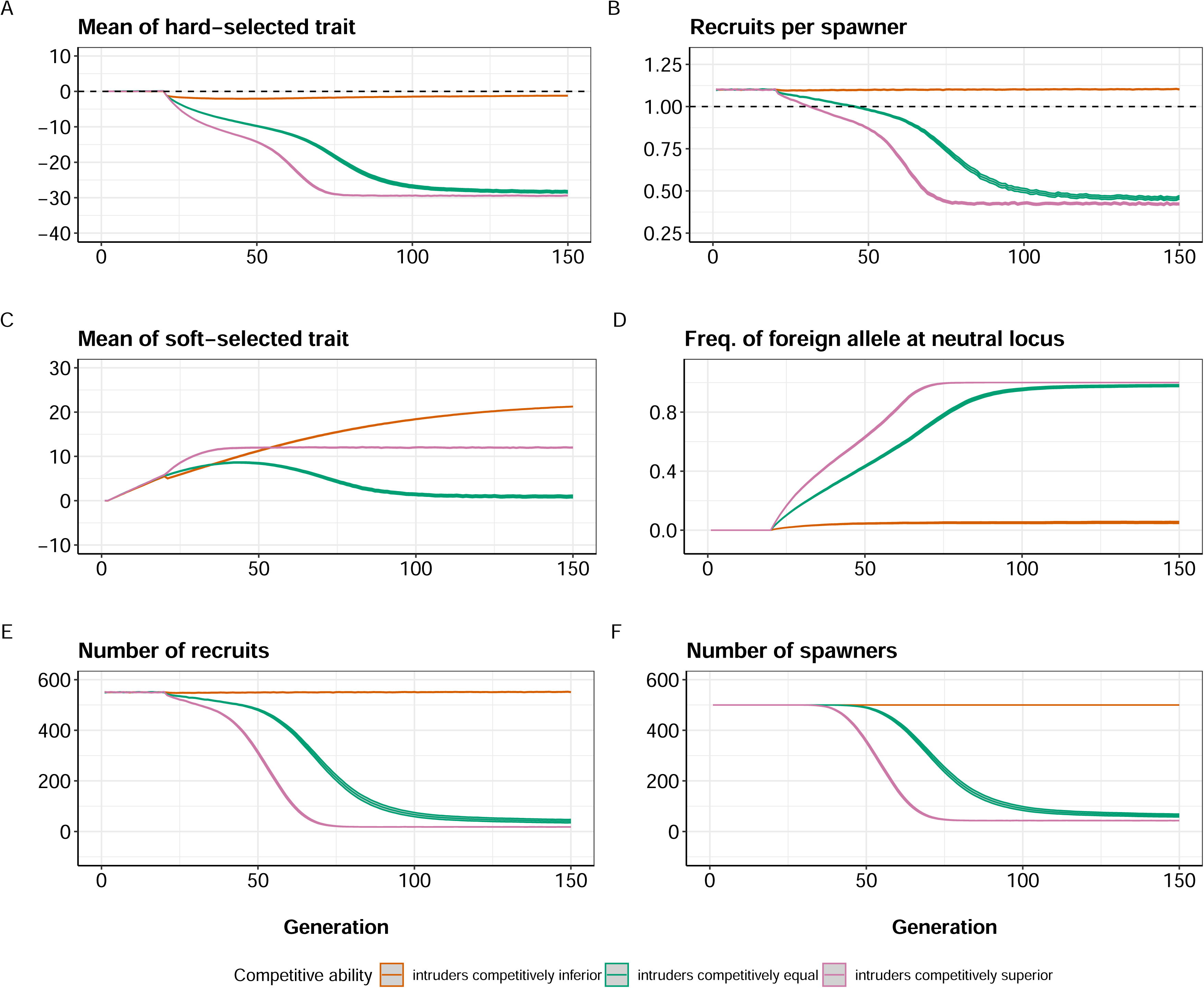
Results of chronic intrusion simulations set 3 for the high reproductive excess scenario (*W*_*MAX*_ = 0.63). Each panel shows the results (mean and 95% confidence intervals across 1000 replicate simulations) comparing cases where the initial heritability (*h^2^*) of both *Z_SOFT_* and *Z_HARD_* was = 0.25 (left sub-panels) or = 0.50 (right sub-panels). The per-generation intrusion rate was fixed at 10% of *K*, i.e., 50 foreign/domesticated fish intruded each generation.

